# Historical biogeography supports Point Conception as the site of turnover between temperate East Pacific ichthyofaunas

**DOI:** 10.1101/2023.05.10.540198

**Authors:** Elizabeth Christina Miller

## Abstract

The northern and southern marine faunas of the temperate Northeastern Pacific meet within California as part of one of the few eastern boundary upwelling ecosystems of the world. Traditionally, it is believed that Point Conception is the precise site of turnover between these two faunas due to sharp changes in oceanographic conditions. However, evidence from intraspecific phylogeography and species range terminals do not support this view, finding stronger biogeographic breaks elsewhere along the coast. Here I develop a new application of historical biogeographic approaches to uncover sites of transition between faunas without needing an *a priori* hypothesis of where these occur. I used this approach to determine whether the point of transition between northern and southern temperate faunas occurs at Point Conception or elsewhere within California. I also take advantage of expert-vetted latitudinal range data of California fish species from the 1970s and the 2020s to assess how biogeography could change with the backdrop of climate change. The site of turnover was found to occur near Point Conception, in concordance with the traditional view. I suggest that recent species- and population-level processes such as dispersal and speciation could be expected to give signals of different events from historical biogeography, possibly explaining the discrepancy across studies. Species richness of California has increased since the 1970s, mostly due to new species from Baja California. Range shifts under warming conditions will have the consequence of increasing the disparity between northern and southern faunas of California, creating a more divergent biogeography.

## INTRODUCTION

A fundamental goal of biogeography is to identify areas with common flora and fauna, and to arrange these areas into a hierarchical classification based on similarity [1,2]. Such classifications are useful to communicate the homology, unity, and evolutionary significance of communities [3]. In general, marine environments pose fewer barriers to dispersal than continental settings [4]. Therefore, marine biogeographic regionalization is often driven by dispersal [5–8]. The high dispersal potential of marine organisms can sometimes work against efforts to classify discrete biogeographic units. When species disperse away from their ancestral lineage’s region of origin, their present-day ranges can give an impression of biogeography that does not reflect history [5].

The politically-demarcated state of California is biogeographically significant because it contains a transition zone where cold-temperate and warm-temperate faunas of the East Pacific meet [9]. Since at least the late 1800s, it has been believed that this transition occurs around Point Conception (34.4°N) [10,11]. At Point Conception, the coastline bends from a north-south to an east-west orientation to form the Southern California Bight. The California Current flows southward from Alaska, running along the coast of North America, until it reaches Point Conception where it is diverted offshore [12]. As a consequence, there is a sharp change in abiotic and biotic conditions within a short distance on either side of Point Conception [13–15]. To the north, temperatures are 2–4° colder, there is strong upwelling and high nutrient concentration, wave action is strong, and the coastline is rocky. Within the Bight, temperatures are warmer, upwelling is weaker, and the coastline is generally sandy and protected from wave action. These factors are believed to pose strong barriers to dispersal that segregate northern and southern faunas [9,12].

Despite these clear environmental differences, the importance of the Point Conception boundary has been controversial for two main reasons. The first reason for controversy comes from the phylogeography literature. Intraspecific genetic breaks for a variety of marine organisms are often not found at Point Conception as expected, and stronger breaks have been found elsewhere along the coast [16–20] (but see [21]). Unlike other well-known features, such as Cape Canaveral in Florida, there is little evidence that Point Conception is preventing gene flow or that it is the site of incipient speciation [16]. Congruent with intraspecific phylogeographic results, northern range terminals of fishes cluster near Los Angeles, further south than Point Conception [9,12]. Overall, there are many more species with ranges that cross Point Conception than species with ranges ending near it [9,22,23]. These results suggest that features other than Point Conception may be more relevant to East Pacific biogeography.

The second reason for controversy is that ranges of the California marine biota are highly dynamic rather than static. The California Current ecosystem is one of only four eastern boundary upwelling systems in the world [24]. The system has undergone regular climate cycles since at least the Pleistocene [25,26]. Examples of these are the Pacific decadal oscillation cycles, which are transitions roughly every 30 years between cool and warm regimes that influence current flow, upwelling, and productivity [27]. In addition, the El Niño southern oscillation occurs on a three-year cycle and similarly alters current action and temperature [26]. These changes incur wholesale northward or southward dispersal of the marine fauna, which are documented in the fossil [28,29] and historical record [30–32]. Hubbs was a vocal critic of attempts to categorize California species into discretized faunas within a hierarchical biogeographic classification, arguing that such an attempt would be thwarted by the inevitability of species’ range shifts [28,33]. More recent classifications have acknowledged this view: a “California Transition Zone” spanning much of the California coastline is recognized by Briggs and Bowen [2] in lieu of a discrete provincial break at Point Conception. This is the only such transition zone recognized in the world’s oceans by their classification, highlighting the unique nature of the California marine fauna.

Both sources of controversy can be potentially ameliorated by historical biogeographical approaches, which consider the role of deep-time events on species’ ranges (Fig. 1). Even if individual species have dispersed away from their parent clade’s center-of-origin, their dispersal routes can be reconstructed based on phylogenetic relationships. For example, rockfishes (*Sebastes*) are one of the most species-rich groups of fishes in California today [34], but they are phylogenetically nested within an otherwise high-latitude clade [35]. This implies that the lineage originally entered California via north-to-south dispersal. This logic can be applied to all members of the California marine fish fauna, because molecular phylogenies sampling all major lineages of fishes have become available in recent years [36,37]. These large phylogenies can be used to investigate the entire marine fish fauna of the East Pacific under a common framework, in order to identify lineages derived from regions north or south of California. The relative proportion of northern-origin to southern-origin species can be overlaid along the coastline of California to reveal transition zones [3,17] without needing an *a priori* hypothesis of where these transitions are located. By incorporating deep-time events (occurring millions of years ago), important areas of transition can be revealed that may otherwise be obfuscated by subsequent dispersal [28] or more recent geologic events driving intraspecific genetic differentiation [17].

**Figure 1.**
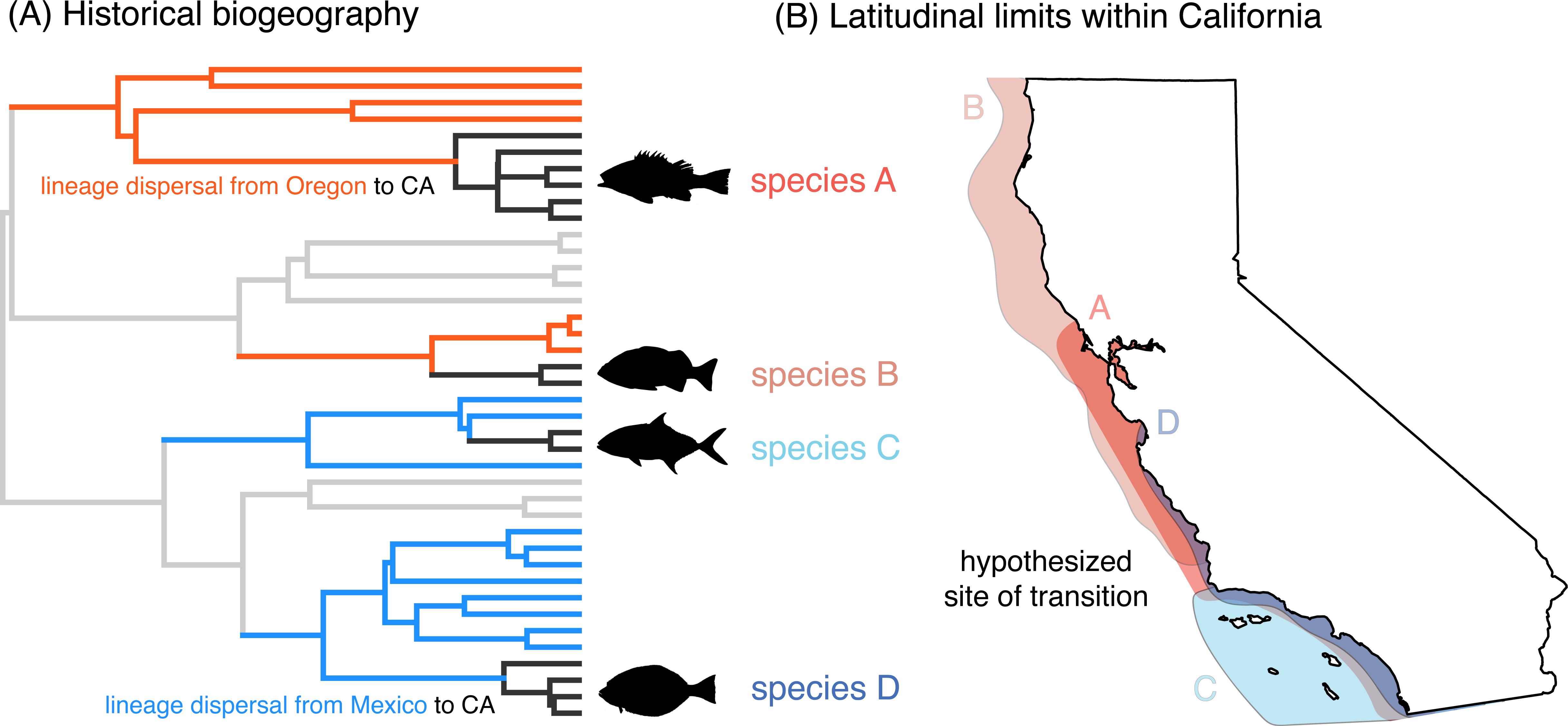
Illustration of the application of historical biogeography for identifying the site of transition between northern and southern faunas within California. (A) The region-of-origin of all California lineages (black) was identified using phylogenetic historical biogeography analyses. Species in California may be derived from southward dispersal (crossing the CA-Oregon border; orange) or northward dispersal (crossing the CA-Mexico border; blue). Light grey branches are outside the temperate East Pacific. Black fish icons are from Phylopic. (B) The latitudinal ranges of species within California were aggregated and compared in light of their region-of-origin (orange=northern origin, blue=southern origin). For example, species A is derived from a northern-origin lineage, but the species’ recent southward dispersal obscures its origins unless the phylogenetic context is considered.

In this study, I utilize probabilistic models of historical biogeography to reconstruct the region-of-origin (either northern or southern affiliation) of California fish species using a phylogenetic context. These models [38–40] reconstruct biogeographic events using present-day ranges of species sampled within a time-calibrated phylogeny. Species’ ranges must be provided in the form of presence or absence within discrete areas of interest. Therefore, these areas must be identified in advance in order to use these methods. In this case, the precise areas of interest are unclear, given the controversy surrounding Point Conception. For this reason, I developed a novel application of these methods to identify regional barriers when they are unknown. I applied this approach to reveal the location of turnover between northern-origin and southern-origin ichthyofaunas along the California coastline.

A secondary goal of this study is to report differences in the biogeography of California potentially driven by anthropogenic climate change. If forecasts are correct, the East Pacific has been in a cold decadal oscillation regime since the late 1990s [27]. Yet, this period has seen several extreme warm events, incurring range shifts of subtropical species into California [32,41,42]. “Ecological surprises” have been documented, such as high anchovy and rockfish recruitment despite warm temperatures, that are suggestive of a novel regime state [43]. The attempt to re-address the biogeography of California in this study is opportunistic of the release of the 2^nd^ edition of Miller and Lea’s seminal guide to California fishes in 2020 [34], which I compared to range estimates from the 1970s [9,44]. This allowed me to construct two impressions of California biogeography based on species ranges during cool and warm periods.

## METHODS

### Data acquisition

I took advantage of expert-vetted sources for the ranges of California marine fishes. Horn and Allen [9] originally published a distributional analysis of marine fishes of California in 1978 (the dataset was obtained through personal communication with L. Allen). Their range data were largely based on the first edition of Miller and Lea [44]. These data were published during a cold decadal oscillation regime [27]. In their study, species were assigned as present or absent within latitudinal bands of 1-degree intervals. For example, a hypothetical species ranging from 34.9°– 32.0° N would be considered present in the 34, 33, and 32-degree bands. There are ten latitudinal bands within California boundaries (from 32–41 degrees). This dataset served as the historical baseline for my study (referred to herein as the 1978 dataset). Note that the authors published an update to their analysis in 2006 [12] with the addition of 22 species known to have dispersed to California from Mexico, but ranges for preexisting species were identical to the 1970s dataset.

The modern dataset (referred to herein as the 2022 dataset) was based on the second edition of the Guide to the Coastal Marine Fishes of California [34] as well as a checklist of species from Alaska to Baja California published in 2021 [45]. These two sources were nearly identical. Following Horn and Allen [9], I assigned species to 1° latitudinal bands to allow backwards compatibility with the historical dataset. This approach has the downside of losing signal at grains finer than 1° (i.e. within latitudinal bands). I used the “validate.names” function in the R package *rfishbase* [46] to convert names to a standard valid name to ensure I was matching up species information correctly across sources. I noted when species’ northern or southern latitudinal limits were different in the recent sources compared to the 1978 dataset (Table S1). I also noted whether species were new to California since the 1978 dataset (Table S2), including species that were described since the original study. A small number of species had unclear range information; these cases are detailed in Table S3. Some of these had to be removed from downstream analyses (Table S3). Human-introduced or invasive species were not considered in this study.

While the original datasets (1978, 2006) recorded latitudinal range boundaries outside of California (i.e., beyond 42.0°N or 32.5°S), I chose to only record limits as “north of CA” or “south of CA” as appropriate. This is because guides were usually not as detailed for ranges outside of California (e.g., stating the northern range boundary as “Alaska”, or southern range boundary as “Peru”, which could span many potential latitudes).

The 1978 study excluded deep-sea fishes. Still, I noted the ranges of deep-sea fishes based on modern sources and included them in biogeographic analyses. While many deep-sea fishes are cosmopolitan, others have clear biogeographic affinities [28,43,47]. Deep-sea fishes were generally defined in this study as those belonging to taxonomic groups that were excluded in the 1978 dataset (such as Myctophiformes or Stomiiformes). Due to high latitude emergence, the usual cut-off of 200 m depth to define a deep-sea species is difficult to apply on a local scale [48]. Biogeographic results for deep-sea fishes are reported separately from inshore fishes (Fig. S1). Only inshore fishes were compared to the 1978 dataset, and reported if new to California waters (Table S2).

### Approach to historical biogeography

I developed a novel application of historical biogeography methods to uncover biogeographic boundaries when their location is uncertain. This approach could be applicable to other biogeographic questions and requires (a) delimiting arbitrary boundaries for a “region” that subsumes putative biogeographic breakpoints of interest, and (b) identifying external regions that represent potential regions-of-origin and dispersal routes. The pattern of boundary-crossing inferred by the ancestral range reconstruction [40] can be used to assign a region-of-origin category to species. The relative proportion of these species categories mapped over a finer spatial scale within the artificial region can then be used to detect patterns of turnover in biogeographic affiliation. In my case, I determined whether species found within California were descended from a lineage that originally entered California via Oregon (USA) or Baja California (Mexico) (Fig. 1). Then, the relative proportion of northern-origin and southern-origin species within each latitudinal band in California was visualized. If Point Conception is the site of turnover of southern-origin to northern-origin communities as expected, then the proportion of these species categories should “flip” from southern-biased to northern-biased beyond the 34° latitudinal band.

Normally it is not advisable to use political boundaries for these models, because historical biogeography pertains to deep-time processes that predate (and are agnostic to) human activities. Therefore, dispersals inferred by the model would refer to lineages crossing an arbitrary boundary that is not biogeographically relevant. In this case, I am purposely setting up artificial boundaries for lineages to cross (one boundary to the north and one to the south), and using this information to infer the direction of dispersal for species within the boundaries. Lineage dispersal and colonization must have happened for species to occur in waters today recognized as Californian, regardless of the political boundary existing or not. Whether lineages crossed the southern or northern artificial boundary is used as evidence to code species as deriving from a northern or southern lineage. These species-specific codes can then be visualized on a finer-grain scale (i.e., 1° latitudinal bands) to visualize patterns of turnover within the arbitrary area of interest, thereby revealing features that *are* biogeographically relevant.

To model the evolution of ranges, I used published time-calibrated molecular phylogenies for Actinopterygii [36] and Chondrichthyes [37]. These phylogenies were constructed from a supermatrix of sequences mined from GenBank. Excellent taxonomic sampling is critical for this approach for two reasons. First, region-of-origin can only be inferred for species sampled in the phylogeny without making tenuous assumptions about biogeography based on taxonomy.

Second, broad sampling of higher taxa is needed to capture dispersal events that occurred in deep time. The ray-finned fish phylogeny used here contains 11,638 species; the cartilaginous fish phylogeny contains 610 species. These phylogenies sampled 73.8% of ray-finned fish and 83.5% of cartilaginous fish species of California. Agnathans were not included in these analyses, but range shifts are still reported in Tables S1 and S2.

Biogeographic transitions were modelled among five “regions”. The temperate East Pacific was broken into three regions: (a) North of the Oregon-California border (equal to the “Cold Temperate Northeast Pacific” of [49] exclusive of California; (b) within California, and (c) South of the California-Mexico border (equal to the “Warm Temperate Northeast Pacific” province exclusive of California). Species were assigned to the “North” and “South” regions based on the checklist by Love et al. [45] (spanning Alaska to Baja California, Mexico). Since the “Warm Temperate Northeast Pacific” province includes the Gulf of California, I also assigned species in this region based on information from the STRI Shorefishes of the Tropical Eastern Pacific database [50]. In total, 784 species were assigned to the “North” region (741 and 43 species of ray-finned and cartilaginous fishes respectively), 902 species were assigned to California (836 and 66), and 1,377 species were assigned to the “South” region (1,270 and 107).

The final two regions were “marine but outside the temperate East Pacific”, and “freshwater”. These regions allowed me to model dispersal into the marine temperate East Pacific itself, as a precursor to dispersal into California. I used a dispersal rate modifier matrix that limited transitions from freshwater into the other four marine states by multiplying the base rate by 0.05. This approach was used in past studies of broad-scale fish biogeography [7,51–53]. The freshwater state was broadly defined to include brackish or diadromous states. Salinity for each species was based on past compilations [51,52] with some updates based on Love and Passarelli [34]. I set the maximum number of areas that a species could co-occur in to five. I removed area combinations from the state space that allowed species to simultaneously occur in the North and South regions but not reside in California. This disjunct state was not observed in any living species. I allowed direct dispersal from the “outside” state into California, but I downweighed the probability of these transitions by 0.05 (as with marine-freshwater transitions). This could be imagined if a lineage crossed the open ocean from the Central Pacific to colonize the East Pacific, which occurs rarely [54].

Biogeographic models were fit using the *BioGeoBEARS* R package v. 1.1.2 [40]. Note that only the BAYAREA class of models [39] are appropriate in this case. This is because these models allow cladogenetic inheritance of ranges spanning multiple regions, which is important because few species are endemic to California itself. In contrast, the DEC class of models [38] would force a widespread range to be broken up into individual regions during cladogenesis, followed by anagenetic dispersal to achieve the widespread range again, inflating the number of dispersals (these issues are discussed in detail in [51]). The fit of the BAYAREA and BAYAREA+J (with founder event speciation) models were compared using Akaike weights. To incorporate uncertainty, I performed biogeographic stochastic mapping [55] to produce 100 independent simulations (“stochastic maps”) of biogeographic history that are possible under the best-fit model. These can be thought of as 100 different boundary-crossing arrangements that can plausibly explain the origins of the California marine ichthyofauna.

To infer the region-of-origin for each California species, I used a custom R script to “walk backwards” from the tip to the branch where dispersal into California occurred (Fig. 1). The source region was obtained for this dispersal event (either the “North” region, “South” region, or “other”). I repeated this sequence for all species and across all 100 stochastic maps. The region-of-origin for a species was considered to be North or South if >70 stochastic maps inferred that region as the source of the ancestral lineage of that species. Of 617 California species, 75.8% could be assigned to a source region this way. Species with the region-of-origin inferred to be North or South in similar frequency across stochastic maps were considered to be of unclear origin (20.2% of species). For the trailing ∼4% of species, the source region was inferred to be “North” plus “other”, or “South” plus “other”, in similar frequencies. Nearly all of these were euryhaline or diadromous species, and the “other” potential source region was “freshwater”. These were assigned to North or South categories respectively.

I generated plots of species richness and north and south range terminals within each of ten latitudinal bands, with species separated by inferred region-of-origin (North, South, or unclear). I also plotted the difference in richness between northern- and southern-origin species within each latitudinal band. I plotted these variables separately using the 1978 and the 2022 range datasets of inshore marine fishes. For additional context, species were coded as demersal or pelagic based on [51] since these life habits can influence dispersal [21].

## RESULTS AND DISCUSSION

### Range shifts of California fishes

The ranges of 151 species of California marine fishes have changed since the 1978 dataset (Table S1). The two most common types of range shifts were: northern range limit extended northward (84 species or 55.6% of range changes); and southern range limit extended southward (43 species or 28.5% of range changes). The majority of range changes involved a species growing its range either northward, southward or in both directions (129 species or 85.4% of range changes).

Two types of range changes are consistent with effects of anthropogenic climate warming: a northward expansion of the northern range limit (caused by dispersal), and a northward expansion of the southern range limit (caused by migration or extirpation). While the former was common, the latter was only seen in 5 species (Table S1). Extensions of the southern range limit further southward were the second most common type. These are not ready explained by climactic changes, since the period 1977–1999 was a warm PDO regime [26,27] which should drive northward dispersal. Range shifts other than the predicted northward direction have been documented in the East Pacific and other marine ecosystems [56,57]. These could be explained by factors such as biotic interactions or idiosyncratic clade-specific responses to warming. Another caveat is that our knowledge of local marine fishes has increased since the 1970s, which could explain some range updates. However, this factor should not drive range changes in any particular direction, whereas shifts driven by warming should generally occur northward.

In addition to range shifts of existing California species, 110 species are new to California waters (Table S2). Of these, 18 species were newly described since the 1978 study, so their occurrence cannot be explained by recent dispersal (Table S2). Among the remaining species, 69 species dispersed from Baja California, Mexico (south-to-north dispersal), 15 species dispersed from Oregon, USA (north-to-south dispersal), with 8 species of unclear origin based on their range (either California endemics or spanning both northern and southern borders).

These counts do not include invasive or introduced species through anthropogenic means, which were discarded from this study.

The greater number of new species added from northward than southward dispersals is consistent with a role of warming climate, either anthropogenic climate change or the recent warm PDO cycle. One complication is that species richness in Mexico is greater than that of Oregon, because richness increases towards the equator (i.e. the latitudinal diversity gradient). Therefore, we would expect the number of northward dispersals to be greater than southward dispersals even under a null model of random dispersal. I performed a post-hoc analysis where I simulated a null distribution of ratios of northward to southward dispersals. The null distribution was made assuming that species have dispersed into California at random since 1978, drawing from the combined pool of species North of California and South of California (details in Note S1). The observed ratio of 4.6 dispersers from Mexico for every 1 disperser from Oregon was well outside the 95% quantile interval of simulated ratios (median simulated ratio=2.5; Fig. S2). These results support the assertion that recent dispersals are influenced by climate warming, even given the confounding influence of the latitudinal diversity gradient.

Due to new species, richness of the California inshore marine fauna has increased 22.3% overall since the 1970s from 494 to 604 species (now including 531 ray-finned fishes, 66 cartilaginous fishes, and 7 jawless fishes). Similar increases in richness primarily due to northward range expansions have been documented in other temperate marine ecosystems [58–60]. Richness increased in all latitudinal bands, but especially in the 32° and 33° bands (including the Mexican border, San Diego, Orange and Los Angeles counties) in which over 75 new species were added to the fauna. This suggests that many of the new species are not dispersing beyond southern California and are affected by the same biogeographic barrier as the pre-1978 fauna (see below).

Like all studies comparing only two time periods, these results should be interpreted with caution [56]. It is unlikely that new species were gradually and consistently added over the 44 years between datasets (1978 and 2022). Many of these new species are known to have entered California during extreme warm events [32,41]. Likewise, it should not be assumed that the historical dataset [9] represents the “normal” or “unmodified” state for California [61]. Fishes in 1978 faced anthropogenic threats as they do today. Cycles of northward and southward dispersals are a persistent feature of the California marine fauna as documented in the fossil and historical record [29–31]; therefore, it is difficult to identify any single snapshot of California’s marine fauna from the past as the “normal” state.

It should be noted that most subtropical species that entered California during the recent warm period from 2014–2018 were not sighted again in 2019 [32]. Whether these new additions will persist in California over the long term is unclear. Breeding populations of at least one new species have been documented [62]. Hubbs [30] noted that subtropical species that entered during the 1853–1860 warm period persisted through the 1880s and were probably breeding within California. With the frequency of extreme warm events predicted to increase in the future [42,63], and given the overall warming trend in California [64], it seems plausible that conditions will eventually remain suitable for subtropical species to establish in California.

### Marine biogeography of California

The BAYAREA+J model had a better fit than the BAYAREA model in both ray-finned fishes and cartilaginous fishes (Akaike weight=1). By “walking backwards” from each tip of the tree to the branch where colonization of California waters occurred, I determined that 188 species of inshore California ray-finned fishes are derived from a northern lineage, 205 species are derived from a southern lineage, and 49 species were of unclear origin (Table S4). Uncertainty was much greater for cartilaginous fishes (2 species from a northern lineage, 29 species from a southern lineage, and 22 species of unclear origin; Table S5). This could be due to the lower richness and smaller size of the cartilaginous fish phylogeny (610 tips versus >11,000 in the ray-finned fish phylogeny) which affects power, and also by the widespread ranges of many species (i.e. spanning most of the East Pacific). Interestingly, while demersal fishes were of northern and southern-origin in similar proportions, pelagic fishes were overwhelmingly of southern origin (Fig. 2). Of deep-sea fishes (Fig. S1), roughly half had unclear affinities, which is unsurprising given their widespread distributions.

**Figure 2.**
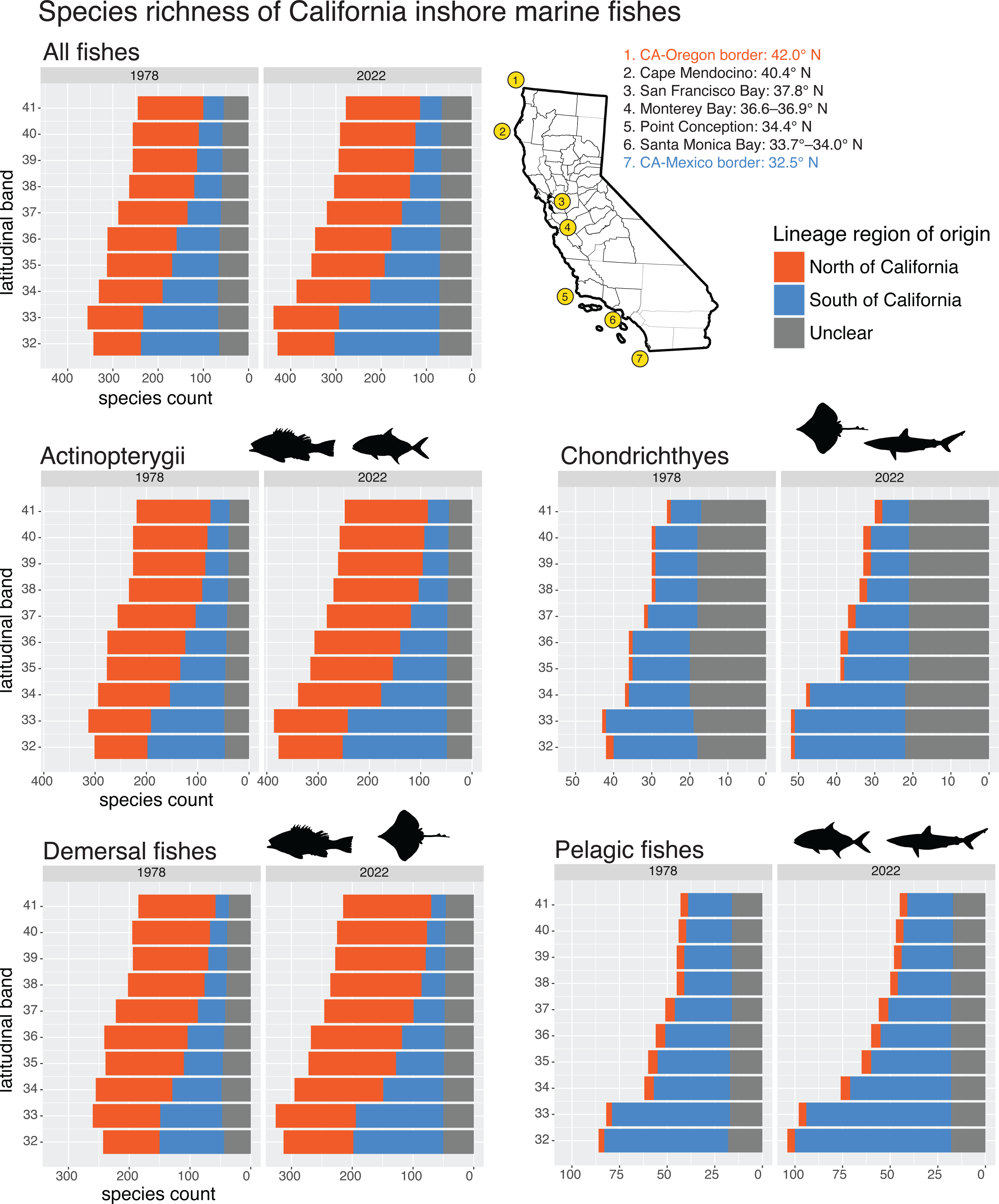
Species richness in each latitudinal band by region-of-origin as inferred from historical biogeography. Range data refer to the 1978 dataset of Horn and Allen [9] or the 2020’s data assembled in this study based on [34,45]. Only inshore marine fishes are shown; for deep-sea fishes see Fig. S1. Region-of-origin for individual species are given in Tables S4 and S5.

Examples of northern-origin lineages include: rockfishes (Sebastidae), sculpins and poachers (Cottoidei), gunnels and pricklebacks (Zoarcoidei), and *Bathyraja* skates (Tables S4, S5). Southern-origin lineages include: many sharks and batoids, surfperches (Embiotocidae), damselfishes (Pomacentridae), scorpionfishes (Scorpaenidae), basses (Serranidae), and jacks (Carangidae). These examples help show how recent speciation and dispersal can erode the signal of events deeper in time. Specifically, surfperches are most diverse today in central and northern California [65], yet they were found to be derived from a southern-origin lineage in my analyses (Table S4). This is because the family Embiotocidae is nested within the larger clade Ovalentaria, which is predominately tropical. Similarly, richness of *Sebastes* rockfishes peaks near Los Angeles in southern California (33° band) [66], yet the lineage was confidently inferred to have colonized California from the northern United States (Table S4). This suggests the modern distribution of rockfishes is due to southward dispersal and speciation within California after their initial colonization. Overall, the novel application of BioGeoBEARS used here helped untangle the influence of deep-time events (colonization of a region by higher taxa) from more recent events (subsequent dispersal of individual species within a region) on the biogeography of California.

Fishes in the 32° and 33° bands were mostly of southern origin (derived from lineages in Baja California) (Figs. 2 and 3). Fishes in the 34° band were of northern- and southern-origins in similar proportions with a slight bias towards northern-origin species. Fishes in the bands from 35°–42° were mostly of northern origin (derived from lineages in Oregon). These patterns were true of both 1978 and 2022 ichthyofaunas. However, the difference between northern and southern California faunas has become more pronounced since the 1970s; that is, southern bands are even more biased towards southern-origin species, and northern bands are even more biased towards northern-origin species, than they were in 1978 (Figs. 2 and 3). This is due to the addition of new species from Baja California (i.e. tropicalization), especially those that do not disperse beyond 33.99° N, with a lesser role from species dispersing southward from northern latitudes (i.e. borealization [57]). Note that deep-sea fishes also showed a south-to-north turnover (Fig. S1), but the transition was more gradual than inshore fishes and there was no obvious biogeographic boundary.

**Figure 3.**
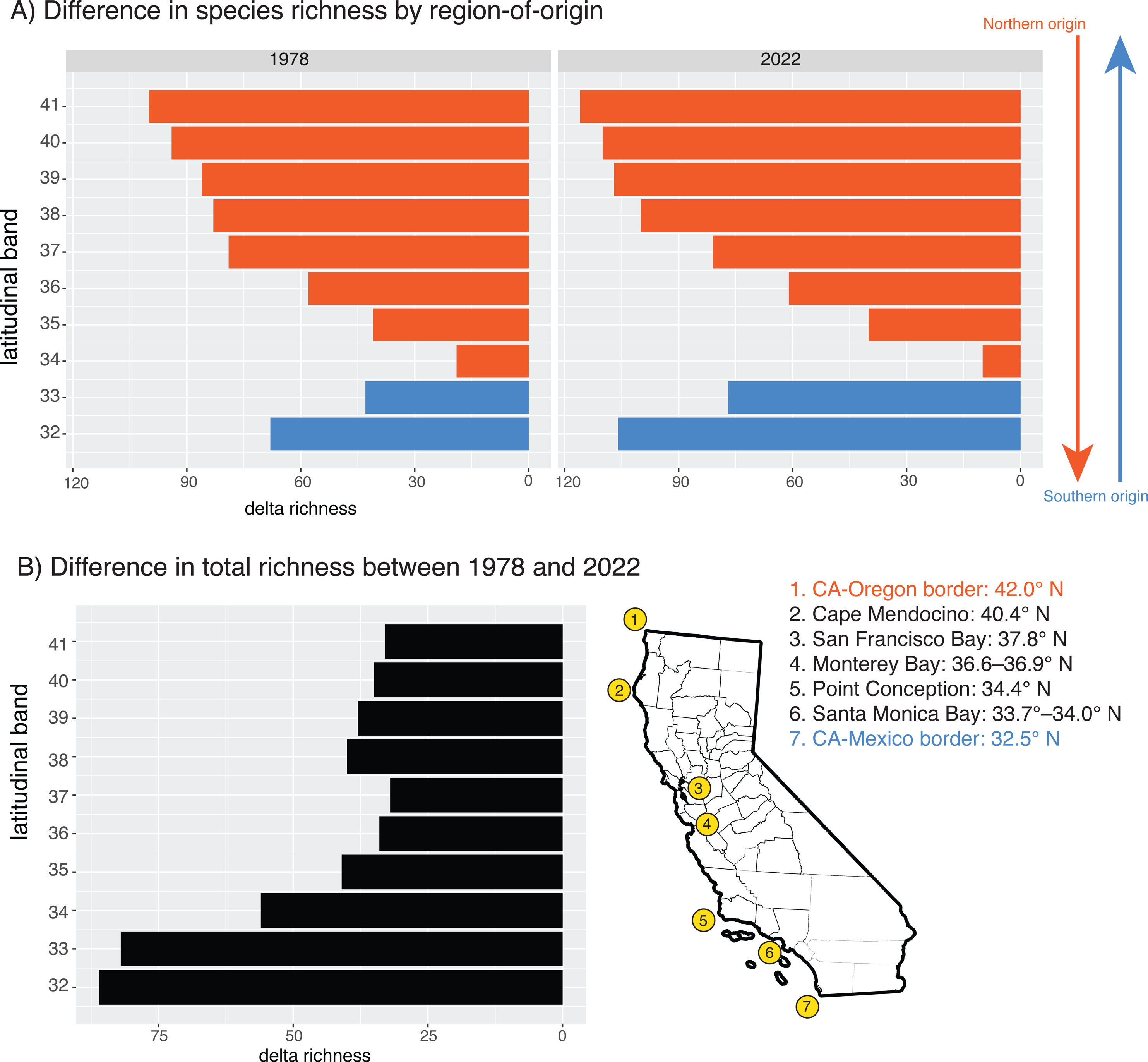
(A) Difference in richness of northern- and southern-origin species within each latitudinal band, using range data from the 1978s and 2020s. Bands are colored by the region-of-origin with the highest richness (blue=southern origin, orange=northern origin). (B) Difference in total species richness in each latitudinal band between the 1978s and 2020s dataset. Only inshore marine species are shown.

If range shifts persist under future warm conditions in California, these results suggest this will have the biogeographic consequence of widening the disparity in faunal composition between communities north and south of Point Conception. It seems that many subtropical species new to California are limited by the same biogeographic barrier as core California species (Figs. 3 and 4). This is unnerving for future survival of species under climate change. If species are prevented from dispersing northward, they may be trapped into unfavorable thermal conditions [67,68]. It is unclear whether continued warming will change oceanographic conditions enough to weaken this barrier in the future.

**Figure 4.**
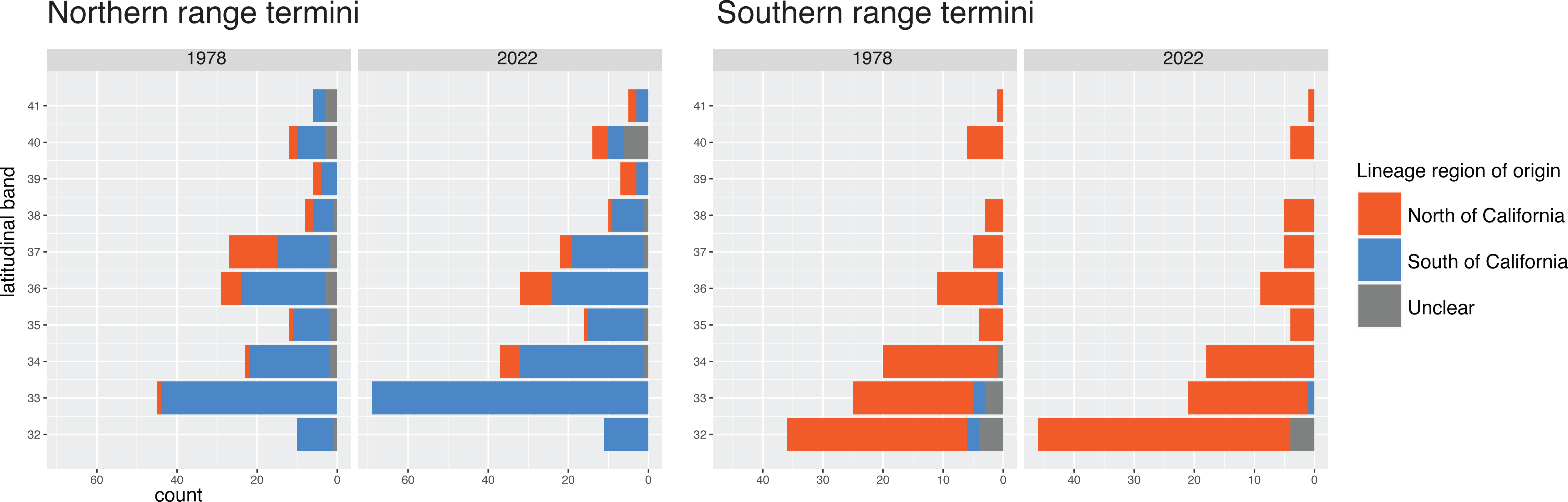
Count of species’ range terminals (northernmost and southernmost records) within each latitudinal band, colored by region-of-origin as inferred from historical biogeography. Only inshore marine species with a range terminal within California are shown.

The locations of northern and southern species range terminals have been traditionally used as evidence for biogeographic boundaries within California [9]. Northern range terminals of southern-origin species peaked in the 33° band (near Los Angeles; Fig. 4). Southern range terminals of northern-origin species peak at 32° N (near San Diego and the Mexican border).

These patterns were also reported by Horn and Allen [9,12] but have become more dramatic in the present day due to range shifts. These results provide support to the suggestion that biogeographic barriers in California are more influential on ranges of southern-origin species than ranges of northern-origin species [9], and this continues to be true for new species entering California. This is consistent with the idea that the southward flowing California Current inhibits recruitment north of Point Conception [21,69].

### Role of Point Conception

Traditionally, it is believed that Point Conception is the barrier with strongest influence on California marine fishes, representing the site of turnover between southern and northern fish faunas of the temperate East Pacific. In favor with this view is the roughly balanced proportion of northern- and southern-origin species within the 34° band (Fig. 3). This band spans 34.0°– 34.99°N, with Point Conception near the center at 34.4° N. The balanced proportions in this band could be explained if southern-origin species outnumber northern-origin species below Point Conception, but the reverse is true above Point Conception. Therefore, the pattern of turnover of northern-origin and southern-origin lineages, as inferred from historical biogeography, is in support of the Point Conception barrier.

Still, an observation against the importance of Point Conception is that southern range terminals concentrate within the 33° band (Fig. 4). If Point Conception was most influential on species’ ranges, we would instead expect southern range terminals to concentrate within the 34° band instead. The observed pattern of range terminals suggests that the most obvious changes in faunal composition occur well outside the vicinity of Point Conception. This is consistent with phylogeographic studies across a variety of marine organisms, which tend to show that intraspecific genetic groupings split near the Los Angeles area [16–18,21,70]. In particular, the site of divergence appears to be Santa Monica Bay (33.7–34.0° N), which is contained by the Palos Verdes peninsula to the south and Point Dume to the north. While Pleistocene sea level change has been implicated as the driver of these genetic splits [17,70], present-day factors must be operating as well, given that many new species entering California during warming events have not dispersed beyond the 33° band (Figs. 3 and 4). One possibility is that sharp transitions in substrate between soft bottoms and rocky reefs and canyons in this area discourages dispersal [70,71].

How do we reconcile the fact that historical biogeography gives a different answer about the relevance of Point Conception than range terminals and intraspecific phylogeography? One answer is that the approach used in this study considers the role of deep-time events on community assembly, specifically the dispersal of ancestral lineages. Subsequent dispersal of individual species away from their center of origin can erode the signal of earlier events [5]. Further, it is worth considering that the California marine fauna is largely dispersal-assembled rather than speciation-assembled. Specifically, faunas on either side of Point Conception are distinct because they are two distantly related groups that have dispersed southward and northward, respectively, and meet in California. This is different from a scenario where the faunas are distinct because they are the product of repeated allopatric speciation events (Burton 1998). The latter scenario may be ongoing in the Los Angeles region, but the conditions that favor genetic structuring in this area may only date to the Pleistocene [17,18,25]. This is long after major fish lineages initially established in California, a process in which Point Conception appears to have been more influential. To sum, these sources of evidence operate on different time scales and can be expected to give signals of different events on biogeographic regionalization.

### Conclusions

Historical biogeographic approaches suggest that the site of turnover of northern-origin to southern-origin ichthyofaunas of the temperate East Pacific occurs near Point Conception, in agreement with the traditional view. This suggests that incorporating deep-time events can help reveal biogeographic boundaries that have been obscured by recent species-level processes. Species richness of California fishes has increased since the 1970s, driven by northward dispersals against the backdrop of climate change. As warming continues, the northern and southern faunas will become more disparate, producing a more divided biogeographic pattern.

## Acknowledgements

I thank Larry Allen, Michael Horn, and Daniel Pondella for providing earlier datasets and for helpful guidance. I thank Ben Frable for suggesting additional sources of data and for advice on the manuscript. Phil Hastings and Dahiana Arcila sponsored a visiting scholar position at time of writing.

## Funding

I was supported by an N.S.F. Postdoctoral Fellowship (DBI-1906574).

## SUPPORTING INFORMATION

All items below can be found in the file **“Supporting Information”:**

Figure S1: Biogeographic affinities of offshore deep-sea fishes by latitudinal band Note S1: Post-hoc analysis of range shifts given the null expectation

Figure S2: Simulated proportion of dispersals from north vs south Table S1: Range shifts of nearshore California fishes since the 1970s

Table S2: New species of nearshore fishes in California waters since the 1970s Table S3: Species with unclear range information

Table S4: Region-of-origin inferred for California ray-finned fish species Table S5: Region-of-origin inferred for California cartilaginous fish species

## Notes

### Competing Interest Statement

The authors have declared no competing interest.

