## Supporting Information for "Historical biogeography supports Point Conception as the site of turnover between temperate East Pacific ichthyofaunas"

Table of Contents:

Figure S1: Biogeographic affinities of offshore deep-sea fishes by latitudinal band

Note S1: Post-hoc analysis of range shifts given the null expectation

Figure S2: Simulated proportion of dispersals from north vs south

Table S1: Range shifts of nearshore California fishes since the 1970s

Table S2: New species of nearshore fishes in California waters since the 1970s

Table S3: Species with unclear range information

Table S4: Region-of-origin inferred for California ray-finned fish species

Table S5: Region-of-origin inferred for California cartilaginous fish species

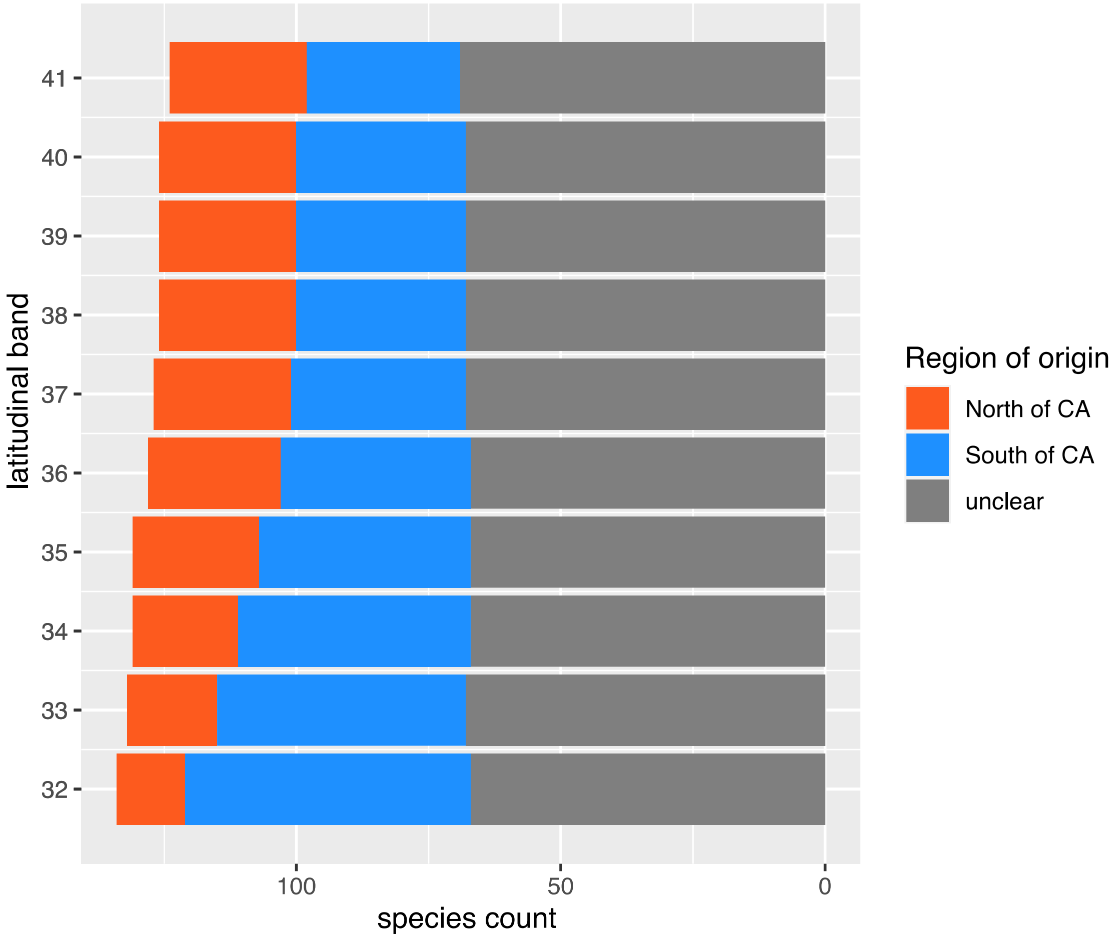

**Figure S1:** Turnover of northern-and-southern origin species with latitudinal band in California waters. This figure shows results for deep-sea species only; see Tables S4–S5 for a list of species included. All range data for these species were based on modern sources (Love and Passarelli 2020; Love et al. 2021).

**Note S1:** Post-hoc analyses of range shifts given the null expectation

In this study I report that most new species in California waters since the 1970s dispersed from Baja California, Mexico (northward dispersal, n=69 species) as opposed to dispersal from Oregon (southward dispersal, n=15 species; Table S2) (excluding new species added from recent species descriptions). This is important because dispersals northward are consistent with an effect of anthropogenic climate change on range shifts. However, this is complicated by the fact that species richness is higher towards the equator (i.e. the latitudinal diversity gradient). Since there are more species in Baja California than in Oregon, the pool of species that can potentially disperse into California is also greater. Therefore, we would expect more species to disperse northward under a null model if dispersal was random.

I simulated potential dispersals to test the hypothesis that the number of northward dispersals since the 1970s was even greater than expected. I determined the pool of species that could have potentially dispersed from Oregon to California since 1978 to be 293 species. This is the number of species known to be restricted to North of California today, plus the number reported to have dispersed to California in this study (these species would have been restricted to North of California in 1978). The same logic was applied to determine the pool of species that could have potentially dispersed from Baja California to California to be 728 species in 1978. I performed trials where I randomly sampled 84 species (the observed number) from the total pool of potential dispersers to disperse to California. I then calculated a ratio of dispersers from the southern pool to dispersers from the northern pool. I performed 100,000 trials to determine a null distribution of ratios.

The median ratio in the simulated distribution is 2.5 dispersal events from Mexico for every 1 dispersal from Oregon. The observed ratio of 4.6 (69 divided by 15 species) is outside of the 95% quantile of the simulated distribution (Figure S2). These results suggest the observed number of species entering California from Mexico is significantly greater than expected at random (given the latitudinal diversity gradient), supporting the assertion that recent dispersals are influenced by anthropogenic climate warming.

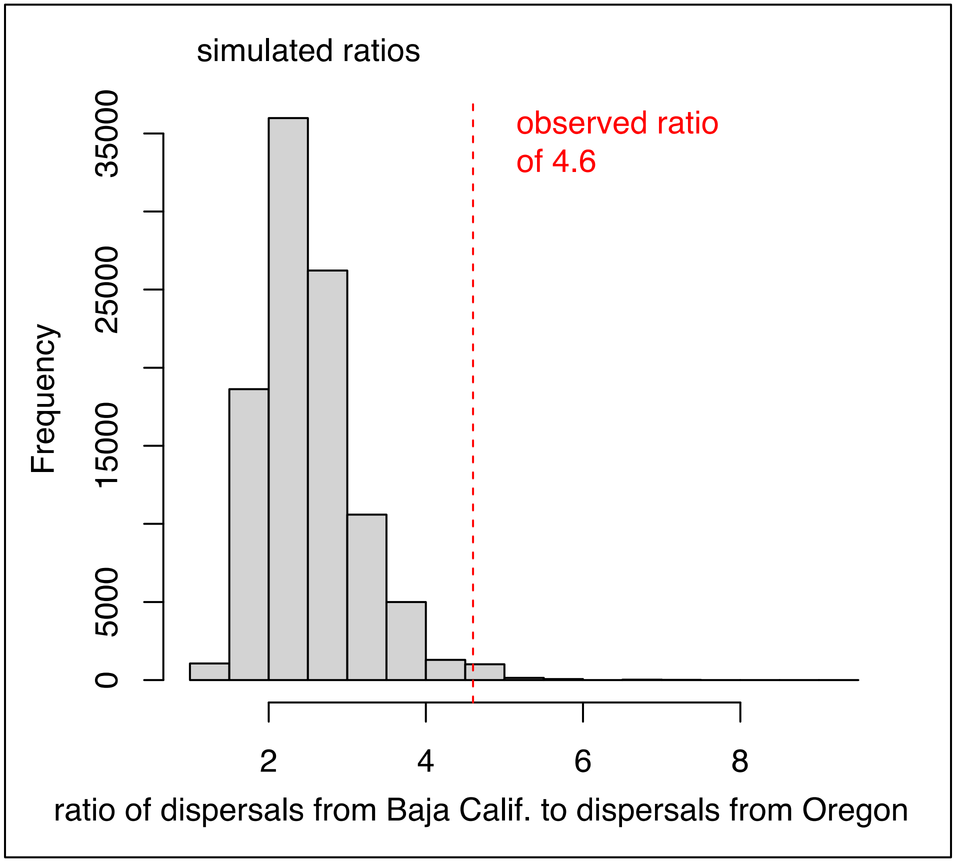

**Figure S2.** Simulated ratios of species dispersing into California from Baja California (Mexico) versus species dispersing from Oregon (USA). The observed ratio is shown by the red dashed line.

**Table S1.** List of range changes among inshore marine species between 1978 dataset (Horn and Allen 1978) and this study.

| Class | Species (other names in databases) | Family | 1970s N limit | New N limit | 1970s S limit | New S limit | Type of change | Sampled in phylogeny |
|---|---|---|---|---|---|---|---|---|
| Actinopterygii | *Artedius corallinus* | Cottidae | N of CA | 41 | S of CA | - | N limit constricted southward | yes |
| Actinopterygii | *Clinocottus recalvus* | Cottidae | 42 | 40 | S of CA | - | N limit constricted southward | yes |
| Actinopterygii | *Lepidocybium flavobrunneum* | Gempylidae | N of CA | 33 | S of CA | - | N limit constricted southward | yes |
| Actinopterygii | *Tetrapturus angustirostris* | Istiophoridae | 42 | 40 | S of CA | - | N limit constricted southward | yes |
| Actinopterygii | *Semicossyphus pulcher* | Labridae | 37 | 36 | S of CA | - | N limit constricted southward | yes |
| Actinopterygii | *Sebastes umbrosus* | Sebastidae | 37 | 36 | S of CA | - | N limit constricted southward | yes |
| Actinopterygii | *Epinephelus analogus* | Serranidae | 34 | 33 | S of CA | - | N limit constricted southward | yes |
| Actinopterygii | *Kathetostoma averruncus* | Uranoscopidae | 39 | 36 | S of CA | - | N limit constricted southward | yes |
| Actinopterygii | *Rathbunella hypoplecta* | Bathymasteridae | N of CA | 34 | S of CA | - | N limit constricted southward (earlier northern records were erroneous) | yes |
| Chondrichthyes | *Dasyatis dipterura (Hypanus dipterurus)* | Dasyatidae | N of CA | 34 | S of CA | - | N limit contracted southward (potentially erroneous in 1970s dataset) | yes |
| Actinopterygii | *Blepsias cirrhosus* | Agonidae | 41 | N of CA | 35 | - | N limit extended northward | yes |
| Actinopterygii | *Leuresthes tenuis* | Atherinopsidae | 37 | 38 | S of CA | - | N limit extended northward | yes |
| Actinopterygii | *Balistes polylepis* | Balistidae | 41 | N of CA | S of CA | - | N limit extended northward | yes |
| Actinopterygii | *Hypsoblennius gilberti* | Blenniidae | 34 | 37 | S of CA | - | N limit extended northward | yes |

| | | | | | | | | |
|---|---|---|---|---|---|---|---|---|
| Actinopterygii | *Hypsoblennius jenkinsi* | Blenniidae | 34 | 36 | S of CA | - | N limit extended northward | yes |
| Actinopterygii | *Pteraclis aesticola* | Bramidae | 35 | 37 | S of CA | - | N limit extended northward | no |
| Actinopterygii | *Taractichthys steindachneri* | Bramidae | 33 | 34 | S of CA | - | N limit extended northward | yes |
| Actinopterygii | *Caranx caninus* | Carangidae | 32 | 33 | S of CA | - | N limit extended northward | yes |
| Actinopterygii | *Naucrates ductor* | Carangidae | 36 | N of CA | S of CA | - | N limit extended northward | yes |
| Actinopterygii | *Trachinotus rhodopus* | Carangidae | 34 | 35 | S of CA | - | N limit extended northward | yes |
| Actinopterygii | *Cebidichthys violaceus* | Cebidichthyidae | 41 | N of CA | S of CA | - | N limit extended northward | yes |
| Actinopterygii | *Neoclinus blanchardi* | Chaenopsidae | 37 | 38 | S of CA | - | N limit extended northward | yes |
| Actinopterygii | *Neoclinus stephensae* | Chaenopsidae | 36 | 37 | S of CA | - | N limit extended northward | no |
| Actinopterygii | *Gibbonsia elegans* | Clinidae | 35 | 37 | S of CA | - | N limit extended northward | yes |
| Actinopterygii | *Clinocottus analis* | Cottidae | 39 | 40 | S of CA | - | N limit extended northward | yes |
| Actinopterygii | *Enophrys taurus (Enophrys taurina)* | Cottidae | 37 | 39 | 33 | - | N limit extended northward | yes |
| Actinopterygii | *Orthonopias triacis* | Cottidae | 37 | N of CA | S of CA | - | N limit extended northward | yes |
| Actinopterygii | *Radulinus vinculus* | Cottidae | 34 | 35 | 34 | - | N limit extended northward | no |
| Actinopterygii | *Symphurus atricauda* | Cynoglossidae | 41 | N of CA | S of CA | - | N limit extended northward | yes |
| Actinopterygii | *Diodon hystrix* | Diodontidae | 32 | 34 | S of CA | - | N limit extended northward | yes |
| Actinopterygii | *Remora remora* | Echeneidae | 37 | N of CA | S of CA | - | N limit extended northward | yes |
| Actinopterygii | *Phanerodon atripes* | Embiotocidae | 38 | N of CA | S of CA | - | N limit extended northward | yes |
| Actinopterygii | *Zalembius rosaceus* | Embiotocidae | 38 | 40 | S of CA | - | N limit extended northward | yes |

| | | | | | | | | |
|---|---|---|---|---|---|---|---|---|
| Actinopterygii | *Cheilopogon heterurus (Cypselurus heterurus)* | Exocoetidae | 32 | 33 | S of CA | - | N limit extended northward | no |
| Actinopterygii | *Ruvettus pretiosus* | Gempylidae | 36 | 38 | S of CA | - | N limit extended northward | yes |
| Actinopterygii | *Eucinostomus dowii* | Gerreidae | 32 | 33 | S of CA | - | N limit extended northward | no |
| Actinopterygii | *Girella nigricans* | Girellidae | 37 | N of CA | S of CA | - | N limit extended northward | yes |
| Actinopterygii | *Rimicola eigenmanni* | Gobiesocidae | 33 | 34 | S of CA | - | N limit extended northward | no |
| Actinopterygii | *Ctenogobius sagittula (Gobionellus longicaudus)* | Gobiidae | 32 | 33 | S of CA | - | N limit extended northward | yes |
| Actinopterygii | *Anisotremus davidsonii* | Haemulidae | 34 | 36 | S of CA | - | N limit extended northward | yes |
| Actinopterygii | *Xenistius californiensis (Haemulon californiensis)* | Haemulidae | 36 | 37 | S of CA | - | N limit extended northward | yes |
| Actinopterygii | *Istiophorus platypterus* | Istiophoridae | 32 | 33 | S of CA | - | N limit extended northward | yes |
| Actinopterygii | *Tetrapturus audax (Kajikia audax)* | Istiophoridae | 34 | N of CA | S of CA | - | N limit extended northward | yes |
| Actinopterygii | *Hermosilla azurea (Kyphosus azureus)* | Kyphosidae | 36 | 41 | S of CA | - | N limit extended northward | yes |
| Actinopterygii | *Halichoeres semicinctus* | Labridae | 34 | 36 | S of CA | - | N limit extended northward | yes |
| Actinopterygii | *Nezumia stelgidolepis* | Macrouridae | 41 | N of CA | S of CA | - | N limit extended northward | no |
| Actinopterygii | *Physiculus rastrelliger* | Moridae | 40 | N of CA | S of CA | - | N limit extended northward | yes |
| Actinopterygii | *Mugil cephalus* | Mugilidae | 36 | 40 | S of CA | - | N limit extended northward | yes |
| Actinopterygii | *Zalieutes elater* | Ogcocephalidae | 34 | N of CA | S of CA | - | N limit extended northward | yes |
| Actinopterygii | *Ophichthus triserialis* | Ophichthidae | 40 | N of CA | S of CA | - | N limit extended northward | yes |
| Actinopterygii | *Lactoria diaphana (Ostracion diaphanum)* | Ostraciidae | 34 | 35 | S of CA | - | N limit extended northward | yes |
| Actinopterygii | *Hippoglossina stomata* | Paralichthyidae | 36 | 37 | S of CA | - | N limit extended northward | yes |

| | | | | | | | | |
|---|---|---|---|---|---|---|---|---|
| Actinopterygii | *Pleuronichthys verticalis* | Pleuronectidae | 37 | N of CA | S of CA | - | N limit extended northward | yes |
| Actinopterygii | *Pristigenys serrula* | Priacanthidae | 34 | N of CA | S of CA | - | N limit extended northward | yes |
| Actinopterygii | *Menticirrhus undulatus* | Sciaenidae | 34 | 35 | S of CA | - | N limit extended northward | yes |
| Actinopterygii | *Auxis rochei* | Scombridae | 33 | 37 | S of CA | - | N limit extended northward | yes |
| Actinopterygii | *Euthynnus affinis* | Scombridae | 33 | 35 | S of CA | - | N limit extended northward | yes |
| Actinopterygii | *Thunnus albacares* | Scombridae | 35 | N of CA | S of CA | - | N limit extended northward | yes |
| Actinopterygii | *Medialuna californiensis (Medialuna californica)* | Scorpididae | 41 | N of CA | S of CA | - | N limit extended northward | yes |
| Actinopterygii | *Sebastes atrovirens* | Sebastidae | 38 | 39 | S of CA | - | N limit extended northward | yes |
| Actinopterygii | *Sebastes chrysomelas* | Sebastidae | 40 | N of CA | S of CA | - | N limit extended northward | yes |
| Actinopterygii | *Sebastes ensifer* | Sebastidae | 37 | N of CA | S of CA | - | N limit extended northward | yes |
| Actinopterygii | *Sebastes eos* | Sebastidae | 37 | N of CA | S of CA | - | N limit extended northward | yes |
| Actinopterygii | *Sebastes gilli* | Sebastidae | 36 | N of CA | 30 | - | N limit extended northward | yes |
| Actinopterygii | *Sebastes hopkinsi* | Sebastidae | 37 | N of CA | S of CA | - | N limit extended northward | yes |
| Actinopterygii | *Sebastes levis* | Sebastidae | 42 | N of CA (northern OR) | S of CA | - | N limit extended northward | no |
| Actinopterygii | *Sebastes ovalis* | Sebastidae | 42 | N of CA (WA state) | S of CA | - | N limit extended northward | yes |
| Actinopterygii | *Sebastes rosenblatti* | Sebastidae | 37 | 40 | S of CA | - | N limit extended northward | yes |
| Actinopterygii | *Sebastes rufus* | Sebastidae | 42 | N of CA (Canada) | S of CA | - | N limit extended northward | yes |
| Actinopterygii | *Sebastes semicinctus* | Sebastidae | 37 | N of CA | S of CA | - | N limit extended northward | yes |

| | | | | | | | | |
|---|---|---|---|---|---|---|---|---|
| Actinopterygii | *Sebastes serranoides* | Sebastidae | 41 | N of CA | S of CA | - | N limit extended northward | yes |
| Actinopterygii | *Epinephelus niphobles (Hyporthodus niphobles)* | Serranidae | 35 | 36 | S of CA | - | N limit extended northward | yes |
| Actinopterygii | *Cosmocampus arctus (Bryx arctos)* | Syngnathidae | 38 | 40 | S of CA | - | N limit extended northward | no |
| Actinopterygii | *Syngnathus californiensis* | Syngnathidae | 37 | 38 | S of CA | - | N limit extended northward | yes |
| Actinopterygii | *Synodus lucioceps* | Synodontidae | 38 | N of CA | S of CA | - | N limit extended northward | yes |
| Actinopterygii | *Sphoeroides annulatus* | Tetraodontidae | 32 | 33 | S of CA | - | N limit extended northward | yes |
| Actinopterygii | *Desmodema lorum (Desmodena polysticta)* | Trachipteridae | 36 | N of CA | S of CA | - | N limit extended northward | no |
| Actinopterygii | *Trachipterus fukuzakii* | Trachipteridae | 33 | 37 | S of CA | - | N limit extended northward | no |
| Actinopterygii | *Zu cristatus* | Trachipteridae | 33 | 34 | S of CA | - | N limit extended northward | yes |
| Actinopterygii | *Lepidopus fitchi* | Trichiuridae | 40 | N of CA | S of CA | - | N limit extended northward | no |
| Actinopterygii | *Trichiurus nitens (Trichiurus lepturus)* | Trichiuridae | 33 | 37 | S of CA | - | N limit extended northward | yes |
| Actinopterygii | *Bellator xenisma* | Triglidae | 33 | 34 | S of CA | - | N limit extended northward | yes |
| Actinopterygii | *Lyconema barbatum* | Zoarcidae | 42 | N of CA (northern Oregon) | S of CA | - | N limit extended northward | no |
| Agnatha | *Myxine circifrons* | Myxinidae | 33 | 37 | S of CA | - | N limit extended northward | NA |
| Chondrichthyes | *Carcharhinus brachyurus (Carcharhinus remotus)* | Carcharhinidae | 33 | 34 | S of CA | - | N limit extended northward | yes |
| Chondrichthyes | *Carcharhinus leucas* | Carcharhinidae | 33 | 34 | S of CA | - | N limit extended northward | yes |
| Chondrichthyes | *Echinorhinus cookei* | Echinorhinidae | 36 | N of CA | S of CA | - | N limit extended northward | yes |
| Chondrichthyes | *Heterodontus francisci* | Heterodontidae | 36 | 37 | S of CA | - | N limit extended northward | yes |
| Chondrichthyes | *Manta birostris (Mobula birostris, Manta hamiltoni)* | Mobulidae | 33 | 34 | S of CA | - | N limit extended northward | yes |

| | | | | | | | | |
|---|---|---|---|---|---|---|---|---|
| Chondrichthyes | *Parmaturus xaniurus* | Pentanchidae | 36 | N of CA | S of CA | - | N limit extended northward | yes |
| Chondrichthyes | *Platyrhinoides triseriata* | Platyrhinidae | 37 | 38 | S of CA | - | N limit extended northward | yes |
| Chondrichthyes | *Rhincodon typus* | Rhincodontidae | 32 | 41 | S of CA | - | N limit extended northward | yes |
| Chondrichthyes | *Mustelus henlei* | Triakidae | 40 | N of CA | S of CA | - | N limit extended northward | yes |
| Chondrichthyes | *Zapteryx exasperata* | Trygonorrhinidae | 33 | 34 | S of CA | - | N limit extended northward | yes |
| Actinopterygii | *Artedius fenestralis* | Cottidae | N of CA | - | 34 | 37 | S limit constricted northward | yes |
| Actinopterygii | *Pseudopentaceros wheeleri (Pentaceros wheeleri)* | Pentacerotidae | N of CA | - | S of CA | 35 | S limit constricted northward | yes |
| Actinopterygii | *Oncorhynchus clarkii* | Salmonidae | N of CA | - | 37 | 40 | S limit constricted northward | yes |
| Actinopterygii | *Oncorhynchus nerka* | Salmonidae | N of CA | - | 33 | 34 | S limit constricted northward | yes |
| Actinopterygii | *Allosmerus elongatus* | Osmeridae | N of CA | - | 33 | 34 | S limit contracted northward | yes |
| Actinopterygii | *Agonus acipenserinus (Podothecus accipenserinus)* | Agonidae | N of CA | - | 40 | 38 | S limit extended southward | yes |
| Actinopterygii | *Asterotheca infraspinata (Bathyagonus infraspinatus)* | Agonidae | N of CA | - | 40 | 38 | S limit extended southward | yes |
| Actinopterygii | *Bathyagonus nigripinnis* | Agonidae | N of CA | - | 40 | 33 | S limit extended southward | yes |
| Actinopterygii | *Bothragonus swanii* | Agonidae | N of CA | - | 35 | 33 | S limit extended southward | no |
| Actinopterygii | *Occella verrucosa (Chesnonia verrucosa)* | Agonidae | N of CA | - | 37 | 34 | S limit extended southward | no |
| Actinopterygii | *Ronquilus jordani* | Bathymasteridae | N of CA | - | 36 | 32 | S limit extended southward | yes |
| Actinopterygii | *Clinocottus acuticeps* | Cottidae | N of CA | - | 36 | 33 | S limit extended southward | yes |
| Actinopterygii | *Enophrys bison* | Cottidae | N of CA | - | 36 | 33 | S limit extended southward | yes |

| | | | | | | | | |
|---|---|---|---|---|---|---|---|---|
| Actinopterygii | *Jordania zonope* | Cottidae | N of CA | - | 34 | 33 | S limit extended southward | yes |
| Actinopterygii | *Oligocottus rimensis* | Cottidae | N of CA | - | 33 | S of CA | S limit extended southward | yes |
| Actinopterygii | *Oligocottus rubellio* | Cottidae | 39 | - | 33 | S of CA | S limit extended southward | yes |
| Actinopterygii | *Radulinus boleoides* | Cottidae | N of CA | - | 33 | 32 (Tanner Bank) | S limit extended southward | yes |
| Actinopterygii | *Amphistichus rhodoterus* | Embiotocidae | N of CA | - | 36 | S of CA | S limit extended southward | yes |
| Actinopterygii | *Gadus macrocephalus* | Gadidae | N of CA | - | 34 | 33 | S limit extended southward | yes |
| Actinopterygii | *Pleurogrammus monopterygius* | Hexagrammidae | N of CA | - | 35 | 33 | S limit extended southward | yes |
| Actinopterygii | *Liparis florae* | Liparidae | N of CA | - | 34 | 33 | S limit extended southward | yes |
| Actinopterygii | *Lumpenus sagitta* | Lumpenidae | N of CA | - | 40 | 36 | S limit extended southward | yes |
| Actinopterygii | *Spirinchus thaleichthys* | Osmeridae | N of CA | - | 37 | 33 | S limit extended southward | yes |
| Actinopterygii | *Thaleichthys pacificus* | Osmeridae | N of CA | - | 38 | 34 | S limit extended southward | yes |
| Actinopterygii | *Apodichthys flavidus* | Pholidae | N of CA | - | 33 | 32 | S limit extended southward | yes |
| Actinopterygii | *Pholis clemensi* | Pholidae | N of CA | - | 38 | 36 | S limit extended southward | yes |
| Actinopterygii | *Pholis ornata* | Pholidae | N of CA | - | 36 | 32 | S limit extended southward | yes |
| Actinopterygii | *Atheresthes stomias* | Pleuronectidae | N of CA | - | 33 | S of CA | S limit extended southward | yes |
| Actinopterygii | *Hippoglossoides elassodon* | Pleuronectidae | N of CA | - | 37 | 32 | S limit extended southward | yes |
| Actinopterygii | *Hippoglossus stenolepis* | Pleuronectidae | N of CA | - | 33 | S of CA | S limit extended southward | yes |
| Actinopterygii | *Isopsetta isolepis* | Pleuronectidae | N of CA | - | 34 | 33 | S limit extended southward | yes |
| Actinopterygii | *Platichthys stellatus* | Pleuronectidae | N of CA | - | 34 | 33 | S limit extended southward | yes |

| | | | | | | | | |
|---|---|---|---|---|---|---|---|---|
| Actinopterygii | *Psettichthys melanostictus* | Pleuronectidae | N of CA | - | 33 | 32 | S limit extended southward | yes |
| Actinopterygii | *Rhamphocottus richardsonii* | Rhamphocottidae | N of CA | - | 33 | 32 | S limit extended southward | yes |
| Actinopterygii | *Sebastes borealis* | Sebastidae | N of CA | - | 36 | 34 | S limit extended southward | yes |
| Actinopterygii | *Sebastes brevispinis* | Sebastidae | N of CA | - | 33 | S of CA | S limit extended southward | yes |
| Actinopterygii | *Sebastes caurinus (Sebastes vexillaris)* | Sebastidae | N of CA | - | 33 | S of CA | S limit extended southward | yes |
| Actinopterygii | *Sebastes melanops* | Sebastidae | N of CA | - | 33 | S of CA | S limit extended southward | yes |
| Actinopterygii | *Sebastes nebulosus* | Sebastidae | N of CA | - | 34 | 33 | S limit extended southward | yes |
| Actinopterygii | *Sebastes nigrocinctus* | Sebastidae | N of CA | - | 35 | 32 | S limit extended southward | yes |
| Actinopterygii | *Anoplarchus purpurescens* | Stichaeidae | N of CA | - | 34 | 33 | S limit extended southward | yes |
| Actinopterygii | *Plagiogrammus hopkinsii* | Stichaeidae | 36 | - | 34 | 33 | S limit extended southward | no |
| Actinopterygii | *Zaprora silenus* | Zaproridae | N of CA | - | 36 | 34 | S limit extended southward | yes |
| Actinopterygii | *Lycodapus mandibularis* | Zoarcidae | N of CA | - | 34 | 32 | S limit extended southward | yes |
| Chondrichthyes | *Somniosus pacificus* | Somniosidae | N of CA | - | 33 | S of CA | S limit extended southward | yes |
| Chondrichthyes | *Lamna ditropis* | Lamnidae | N of CA | - | 32 | S of CA | S limit extended southward (central Baja) | yes |
| Chondrichthyes | *Raja stellulata (Beringraja stellulata)* | Rajidae | N of CA | - | 32 | S of CA | S limit extended southward (central Baja) | yes |
| Chondrichthyes | *Bathyraja abyssicola* | Arhynchobatidae | N of CA | - | 32 | S of CA | S limit extended southward (Galapagos Is) | yes |
| Actinopterygii | *Lophotus capellei (Lophotus cristatus)* | Lophotidae | 32 | 38 | S of CA | 32 (San Diego) | Both limits shifted northward (range in 1970s dataset may be erroneous) | yes |
| Actinopterygii | *Eucyclogobius newberryi* | Gobiidae | N of CA | 41 | 32 | 33 | Both N and S limits constricted | yes |

| | | | | | | | | |
|---|---|---|---|---|---|---|---|---|
| Actinopterygii | *Icelinus fimbriatus* | Cottidae | 36 | N of CA | 33 | 32 | Both N and S limits extended | yes |
| Actinopterygii | *Sebastes lentiginosus* | Sebastidae | 33 | 34 | 32 | S of CA (southern Baja) | Both N and S limits extended | yes |
| Actinopterygii | *Sebastes phillipsi* | Sebastidae | 37 | 41 | 33 | 32 | Both N and S limits extended | yes |
| Actinopterygii | *Hippocampus ingens* | Syngnathidae | 33 | 34 | S of CA | 32 (San Diego) | Both N and S limits shifted northward | yes |
| Chondrichthyes | *Alopias superciliosus* | Alopiidae | N of CA | 40 | 33 | S of CA | Both N and S limits shifted southward | yes |
| Chondrichthyes | *Euprotomicrus bispinatus* | Dalatiidae | 40 | 35 | 33 | S of CA | Both N and S limits shifted southward | yes |
| Chondrichthyes | *Chlamydoselachus anguineus* | Chlamydoselachidae | N of CA | 34 | 34 | S of CA | Range from 70s dataset seems erroneous | yes |

**Table S2.** List of new inshore marine species added to California waters since 1978 dataset (Horn and Allen 1978). Ref. [1] = Love and Passarelli 2020. Ref. [2] = Love et al. 2021. Ref. [3] = Horn et al. 2006. Species with a (*) were newly described or revised since the 1978 study, so their addition cannot be attributed to dispersal (see Love and Passarelli 2020 and references therein).

| Class | Species (other names in databases) | Family | In 2006 dataset | N limit | S limit | Sampled in phylogeny | Notes |
|---|---|---|---|---|---|---|---|
| Actinopterygii | *Acanthurus xanthopterus* | Acanthuridae | no | 33 | S of CA | yes | |
| Actinopterygii | *Agonomalus mozinoi** | Agonidae | no | N of CA | 35 | no | |
| Actinopterygii | *Anoplagonus inermis* | Agonidae | no | N of CA | 38 | yes | |
| Actinopterygii | *Bathyagonus alascanus* | Agonidae | no | N of CA | 40 | yes | |
| Actinopterygii | *Hemitripterus bolini* | Agonidae | no | N of CA | 40 | no | |
| Actinopterygii | *Pallasina aix (Pallasina barbata)* | Agonidae | no | N of CA | 37 | no | |
| Actinopterygii | *Apogon atricaudus* | Apogonidae | no | 33 | S of CA | no | |
| Actinopterygii | *Apogon retrosella* | Apogonidae | no | 32 | S of CA | no | |
| Actinopterygii | *Rathbunella alleni* | Bathymasteridae | no | N of CA | S of CA | no | |
| Actinopterygii | *Ophioblennius steindachneri* | Blenniidae | no | 33 | S of CA | yes | |
| Actinopterygii | *Selar crumenophthalmus* | Carangidae | no | 33 | S of CA | yes | |
| Actinopterygii | *Chanos chanos* | Chanidae | no | 33 | S of CA | yes | |
| Actinopterygii | *Opisthonema libertate* | Clupeidae | no | 34 | S of CA | yes | |
| Actinopterygii | *Opisthonema medirastre* | Clupeidae | no | 33 | S of CA | yes | |
| Actinopterygii | *Coryphaena equiselis* | Coryphaenidae | no | 32 | S of CA | yes | |
| Actinopterygii | *Asemichthys taylori* | Cottidae | no | N of CA | 36 | no | |
| Actinopterygii | *Cottus asper* | Cottidae | no | N of CA | 34 | yes | |
| Actinopterygii | *Icelinus limbaughi** | Cottidae | no | 34 | 32 | no | |
| Actinopterygii | *Symphurus oligomerus** | Cynoglossidae | no | 34 | S of CA | no | |
| Actinopterygii | *Diodon eydouxii* | Diodontidae | no | 33 | S of CA | no | |
| Actinopterygii | *Fistularia commersonii* | Fistulariidae | no | 33 | S of CA | yes | |
| Actinopterygii | *Bollmannia stigmatura* | Gobiidae | no | 33 | S of CA | no | |
| Actinopterygii | *Eucyclogobius kristinae** | Gobiidae | no | 33 | 32 | no | |
| Actinopterygii | *Conodon serrifer* | Haemulidae | no | 33 | S of CA | yes | |
| Actinopterygii | *Haemulon flaviguttatum* | Haemulidae | no | 33 | S of CA | yes | |
| Actinopterygii | *Haemulopsis axillaris* | Haemulidae | no | 32 | S of CA | yes | |
| Actinopterygii | *Haemulopsis elongatus* | Haemulidae | no | 32 | S of CA | yes | |
| Actinopterygii | *Microlepidotus inornatus* | Haemulidae | no | 33 | S of CA | yes | |
| Actinopterygii | *Hexagrammos stelleri* | Hexagrammidae | no | N of CA | 37 | yes | |
| Actinopterygii | *Sectator ocyurus (Kyphosus ocyurus)* | Kyphosidae | no | 35 | S of CA | no | |

| | | | | | | | |
|---|---|---|---|---|---|---|---|
| Actinopterygii | *Labrisomus xanti* | Labrisomidae | no | 34 | S of CA | yes | |
| Actinopterygii | *Caulolatilus affinis* | Latilidae | no | 34 | S of CA | yes | |
| Actinopterygii | *Liparis nanus (Lipariscus nanus)* | Liparidae | no | N of CA | 36 | yes | |
| Actinopterygii | *Lophiodes caulinaris* | Lophiidae | no | 37 | S of CA | yes | |
| Actinopterygii | *Lophiodes spilurus* | Lophiidae | no | 36 | S of CA | yes | |
| Actinopterygii | *Lutjanus argentiventris* | Lutjanidae | no | 33 | S of CA | yes | |
| Actinopterygii | *Lutjanus colorado* | Lutjanidae | no | 35 | S of CA | yes | |
| Actinopterygii | *Lutjanus novemfasciatus* | Lutjanidae | no | 35 | S of CA | yes | |
| Actinopterygii | *Lutjanus peru* | Lutjanidae | no | 32 | S of CA | yes | |
| Actinopterygii | *Mugil curema* | Mugilidae | no | 34 | S of CA | yes | |
| Actinopterygii | *Muraena argus* | Muraenidae | no | 33 | S of CA | yes | |
| Actinopterygii | *Brotula clarkae* | Ophidiidae | no | 33 | S of CA | no | |
| Actinopterygii | *Kasatkia seigeli** | Opisthocentridae | no | 39 | 35 | yes | |
| Actinopterygii | *Inopsetta ischyra* | Pleuronectidae | no | N of CA | 37 | no | hybrid species of *Pleuronectus vetulus* and *Platichthys stellatus* |
| Actinopterygii | *Pomacanthus zonipectus* | Pomacanthidae | no | 33 | S of CA | yes | |
| Actinopterygii | *Abudefduf troschelii* | Pomacentridae | no | 33 | S of CA | yes | |
| Actinopterygii | *Azurina hirundo* | Pomacentridae | no | 33 | S of CA | yes | |
| Actinopterygii | *Chromis alta** | Pomacentridae | no | 33 | S of CA | yes | |
| Actinopterygii | *Stegastes leucorus* | Pomacentridae | no | 33 | S of CA | no | |
| Actinopterygii | *Heteropriacanthus carolinus (Heteropriacanthus cruentatus)* | Priacanthidae | no | 33 | S of CA | yes | |
| Actinopterygii | *Acanthocybium solandri* | Scombridae | no | 34 | S of CA | yes | |
| Actinopterygii | *Scorpaena mystes* | Scorpaenidae | no | 33 | S of CA | yes | |
| Actinopterygii | *Sebastes diaconus** | Sebastidae | no | N of CA | 34 | no | |
| Actinopterygii | *Sebastes emphaeus* | Sebastidae | no | N of CA | 34 | yes | |
| Actinopterygii | *Sebastes melanosema** | Sebastidae | no | 34 | S of CA | yes | |
| Actinopterygii | *Sebastes melanostictus* | Sebastidae | no | N of CA | 32 | yes | |
| Actinopterygii | *Sebastes moseri** | Sebastidae | no | 36 | S of CA | yes | |
| Actinopterygii | *Sebastes rufinanus** | Sebastidae | no | 34 | S of CA | yes | |
| Actinopterygii | *Cephalopholis colonus (Paranthias colonus)* | Serranidae | no | 32 | S of CA | yes | |
| Actinopterygii | *Dermatolepis dermatolepis* | Serranidae | no | 33 | S of CA | yes | |
| Actinopterygii | *Epinephelus labriformis* | Serranidae | no | 32 | S of CA | yes | |
| Actinopterygii | *Hyporthodus acanthistius* | Serranidae | no | 34 | S of CA | yes | |
| Actinopterygii | *Paralabrax auroguttatus* | Serranidae | no | 34 | S of CA | yes | |

| | | | | | | | |
|---|---|---|---|---|---|---|---|
| Actinopterygii | *Pronotogrammus multifasciatus (Anthias gordensis)* | Serranidae | no | 34 | S of CA | yes | |
| Actinopterygii | *Serranus aequidens* | Serranidae | no | 33 | S of CA | no | |
| Actinopterygii | *Esselenichthys carli** | Stichaeidae | no | 36 | S of CA | yes | |
| Actinopterygii | *Esselenichthys laurae** | Stichaeidae | no | 37 | S of CA | no | |
| Actinopterygii | *Lumpenopsis clitella** | Stichaeidae | no | 33 | 32 | no | |
| Actinopterygii | *Syngnathus euchrous** | Syngnathidae | no | 33 | S of CA | no | |
| Actinopterygii | *Syngnathus exilis* | Syngnathidae | no | 37 | S of CA | yes | |
| Actinopterygii | *Prionotus albirostris* | Triglidae | no | 32 | S of CA | no | |
| Actinopterygii | *Astroscopus zephyreus* | Uranoscopidae | no | 33 | S of CA | no | |
| Actinopterygii | *Eucryphycus californicus* | Zoarcidae | no | 36 | 32 | yes | |
| Actinopterygii | *Lycodes brevipes* | Zoarcidae | no | N of CA | 34 | yes | |
| Agnatha | *Eptatretus mcconnaugheyi** | Myxinidae | no | 33 | S of CA | NA | |
| Agnatha | *Myxine hubbsi* | Myxinidae | no | unclear | S of CA | NA | [1]: "Southern CA" to Chile |
| Chondrichthyes | *Alopias pelagicus* | Alopiidae | no | unclear | S of CA | yes | [1] give northern limit as "southern CA" |
| Chondrichthyes | *Bathyraja aleutica* | Arhynchobatidae | no | N of CA | 36 | yes | |
| Chondrichthyes | *Nasolamia velox* | Carcharhinidae | no | 34 | S of CA | yes | |
| Chondrichthyes | *Hydrolagus melanophasma** | Chimaeridae | no | 36 | S of CA | yes | |
| Chondrichthyes | *Isistius brasiliensis* | Dalatiidae | no | 37 | S of CA | yes | |
| Chondrichthyes | *Centroscyllium nigrum* | Etmopteridae | no | unclear | S of CA | yes | [1] gives northern limit as "southern California" |
| Chondrichthyes | *Isurus paucus* | Lamnidae | no | unclear | S of CA | yes | [1]: Only two records known from CA waters, range is unclear within California |
| Chondrichthyes | *Megachasma pelagios* | Megachasmidae | no | unclear | S of CA | yes | [1]: "occasional off CA". Range within California is unclear |
| Chondrichthyes | *Apristurus kampae** | Pentanchidae | no | N of CA | S of CA | yes | |
| Chondrichthyes | *Amblyraja hyperborea* | Rajidae | no | N of CA | S of CA | yes | |
| Chondrichthyes | *Zameus squamulosus* | Somniosidae | no | 33 | S of CA | yes | |
| Chondrichthyes | *Sphyrna lewini* | Sphyrnidae | no | 34 | S of CA | yes | |
| Actinopterygii | *Apogon pacificus* | Apogonidae | yes | 33 | S of CA | yes | Range extended further northward |

| | | | | | | | |
|---|---|---|---|---|---|---|---|
| | | | | | | | from 32 to 33 degree band since [3] |
| Actinopterygii | *Plagiotremus azaleus* | Blenniidae | yes | 33 | S of CA | yes | |
| Actinopterygii | *Engyophrys sanctilaurentii* | Bothidae | yes | 32 | S of CA | no | Range corrected from 33 since [3] (potentially erroneous) to 32 [1] |
| Actinopterygii | *Synchiropus atrilabiatus* | Callionymidae | yes | 33 | S of CA | yes | |
| Actinopterygii | *Caranx sexfasciatus* | Carangidae | yes | 32 | S of CA | yes | |
| Actinopterygii | *Caranx vinctus* | Carangidae | yes | 32 | S of CA | yes | |
| Actinopterygii | *Selene brevoortii* | Carangidae | yes | 33 | S of CA | yes | Range extended further northward from 32 to 33 degree band since [3] |
| Actinopterygii | *Ruscarius creaseri* | Cottidae | yes | 36 | S of CA | yes | |
| Actinopterygii | *Citharichthys fragilis* | Cyclopsettidae | yes | 34 | S of CA | no | Range extended further northward from 33 to 34 degree band since [3] |
| Actinopterygii | *Diodon holocanthus* | Diodontidae | yes | 37 | S of CA | yes | Range extended further northward from 32 to 37 degree band since [3] |
| Actinopterygii | *Elops affinis* | Elopidae | yes | 34 | S of CA | yes | |
| Actinopterygii | *Fistularia corneta* | Fistulariidae | yes | 33 | S of CA | yes | |
| Actinopterygii | *Rimicola cabrilloi** | Gobiesocidae | yes | 34 | 33 | no | |
| Actinopterygii | *Kyphosus analogus (Kyphosus vaigiensis)* | Kyphosidae | yes | 36 | S of CA | yes | Range extended further northward from 33 to 36 degree band since [3] |
| Actinopterygii | *Decodon melasma* | Labridae | yes | 33 | S of CA | yes | |
| Actinopterygii | *Nicholsina denticulata* | Labridae | yes | 33 | S of CA | yes | |
| Actinopterygii | *Lobotes pacifica* | Lobotidae | yes | 33 | S of CA | yes | |
| Actinopterygii | *Diplectrum maximum* | Serranidae | yes | 33 | S of CA | yes | |

| | | | | | | | |
|---|---|---|---|---|---|---|---|
| Actinopterygii | *Sphyraena ensis* | Sphyraenidae | yes | 33 | S of CA | yes | |
| Actinopterygii | *Anoplarchus insignis* | Stichaeidae | yes | N of CA | 38 | yes | |
| Actinopterygii | *Chirolophis decoratus* | Stichaeidae | yes | N of CA | 37 | yes | Range extended further southward from 40 to 37 degree band since [3] |
| Actinopterygii | *Sphoeroides lobatus* | Tetraodontidae | yes | 33 | S of CA | yes | |

**Table S3:** Species with ambiguous information that are excluded from some analyses. Ref. [1] = Love and Passarelli 2020. Ref. [2] = Love et al. 2021. Ref. [3] = Horn et al. 2006.

| Class | Species (other names in databases) | Family | 1970s N limit | 1970s S limit | Justification | Used in BioGeoBEARS | Change in range in 2020s? |
|---|---|---|---|---|---|---|---|
| Actinopterygii | *Ammodytes hexapterus* | Ammodytidae | N of CA | 33 | [2]: earlier California records referring to *A. hexapterus* are actually records for *A. personatus*; *A. hexapterus* is now a separate species restricted to N of CA. | Yes (under *A. personatus*) | No (under *A. personatus* in recent texts) |
| Actinopterygii | *Liparis rutteri* | Liparidae | N of CA | 33 | [2]: This species has been split; *L. rutteri* is no longer in CA waters. Data from 1970s refers to species now called *L. adiastolus*. *L. rutteri* is in the phylogeny, but treated as *L. adiastolus* in analyses | Yes (under outdated name *Liparis rutteri*; data for *L. adiastolus*) | No |
| Actinopterygii | *Melichthys niger* | Balistidae | 32 | S of CA | [2] claims the San Diego record dates to the 1800's, and the species has not been seen in CA waters since. | Yes (removed record from CA) | - |
| Actinopterygii | *Brama dussumieri* | Bramidae | - | - | [2]: only one record in California, normally a Japanese species | Excluded- cannot determine whether resident in California | Not found in 1970s dataset |
| Actinopterygii | *Oligoplites saurus* | Carangidae | 33 | S of CA | [2] contest that California records are erroneous, even though it was included in 1970's dataset | yes | Removed from CA (corrected error) |
| Actinopterygii | *Cetichthys parini* | Cetomimidae | - | - | [2]: only two records in California | Excluded- too little information to determine range | Not found in 1970s dataset (deep-sea fish) |

| | | | | | | | |
|---|---|---|---|---|---|---|---|
| Actinopterygii | *Parataeniophorus brevis* | Cetomimidae | - | - | [2] give range as "southern CA" | Excluded- cannot determine whether range extends beyond California | Not found in 1970s dataset (deep-sea fish) |
| Actinopterygii | *Kali falx* | Chiasmodontidae | - | - | [2]: "in northern hemisphere". Not specific enough | Excluded- too little information to determine range | Not found in 1970s dataset (deep-sea fish) |
| Actinopterygii | *Cottus aleuticus* | Cottidae | - | - | Sometimes found in nearshore marine waters [1] but its range in marine habitats within CA is unclear | yes | Not in 1970s dataset (mostly freshwater) |
| Actinopterygii | *Gasterosteus aculeatus* | Gasterosteidae | N of CA | S of CA | It appears that the 1970s dataset used its freshwater range, which is larger than its marine range (extends to S of CA). I prefer to use its marine range instead (S limit 36), and due to discrepancies it was removed from range change plot | Yes (coded using marine range) | Unclear |
| Actinopterygii | *Gigantactis gargantua* | Gigantactinidae | - | - | [2] give range as "southern CA" | Excluded- cannot determine whether range extends beyond California | Not found in 1970s dataset (deep-sea fish) |
| Actinopterygii | *Lampris guttatus* | Lampridae | N of CA | S of CA | This is no longer a valid species (see [1] and [2]) but still represented in the phylogeny. Included in BioGeoBEARS but not in range change plots | Yes (under outdated name *Lampris guttatus*) | No longer a valid species |
| Actinopterygii | *Linophryne racemifera* | Linophrynidae | - | - | [2] give range as "southern CA" | Excluded- cannot determine whether range extends beyond California | Not found in 1970s dataset (deep-sea fish) |

| | | | | | | | |
|---|---|---|---|---|---|---|---|
| Actinopterygii | *Centrobranchus nigroocellatus* | Myctophidae | - | - | [2]: only one record in California | Excluded- too little information to determine range | Not found in 1970s dataset (deep-sea fish) |
| Actinopterygii | *Diaphus fulgens* | Myctophidae | - | - | [2]: range poorly known | Excluded- too little information to determine range | Not found in 1970s dataset (deep-sea fish) |
| Actinopterygii | *Diaphus kuroshio* | Myctophidae | - | - | [2]: only one record in California | Excluded- too little information to determine range | Not found in 1970s dataset (deep-sea fish) |
| Actinopterygii | *Diaphus parri* | Myctophidae | - | - | [2]: only one record in California | Excluded- too little information to determine range | Not found in 1970s dataset (deep-sea fish) |
| Actinopterygii | *Diaphus rafinesquii* | Myctophidae | - | - | [2]: range poorly known | Excluded- too little information to determine range | Not found in 1970s dataset (deep-sea fish) |
| Actinopterygii | *Diaphus trachops* | Myctophidae | - | - | [2] give range as “central CA” | Excluded- cannot determine whether range extends beyond California | Not found in 1970s dataset (deep-sea fish) |
| Actinopterygii | *Lampanyctus nobilis* | Myctophidae | - | - | [2]: only one record in California | Excluded- too little information to determine range | Not found in 1970s dataset (deep-sea fish) |
| Actinopterygii | *Notoscopelus japonicus* | Myctophidae | - | - | [2]: unconfirmed west of CA. | Excluded- too little information to determine range | Not found in 1970s dataset (deep-sea fish) |
| Actinopterygii | *Dolopichthys longicornis* | Oneirodidae | - | - | [2] give range as “northern CA” | Excluded- cannot determine whether range extends beyond California | Not found in 1970s dataset (deep-sea fish) |

| | | | | | | | |
|---|---|---|---|---|---|---|---|
| Actinopterygii | *Microlophichthys microlophus* | Oneirodidae | - | - | [2] give range as "well off southern CA" | Excluded- too little information to determine range | Not found in 1970s dataset (deep-sea fish) |
| Actinopterygii | *Bassozetus levistomatus* | Ophidiidae | - | - | [2]: only one record in California | Excluded- too little information to determine range | Not found in 1970s dataset |
| Actinopterygii | *Ioichthys kashkini* | Opisthoproctidae | - | - | [2]: only one record in California | Excluded- too little information to determine range | Not found in 1970s dataset (deep-sea fish) |
| Actinopterygii | *Oplegnathus fasciatus* | Oplegnathidae | - | - | [2]: only entered CA as a vagrant | Excluded- not a resident in CA | Not found in 1970s dataset |
| Actinopterygii | *Hypomesus transpacificus* | Osmeridae | - | - | Mentioned in [1] but is predominately a freshwater species | Excluded- nonmarine | Not found in 1970s dataset (mostly freshwater) |
| Actinopterygii | *Allothunnus fallai* | Scombridae | 33 | S of CA | [2]: two isolated reports in the focal region. Seems to be vagrant in California (even though was included in 1970s dataset) | Excluded- not a resident in CA | - |
| Actinopterygii | *Sebastes aleutianus* | Sebastidae | N of CA | 32 | [2]: southern limit is northern California. [2] say this species was recently recognized, and its range limits are unclear due to confusion with other species | yes | Unclear |
| Actinopterygii | *Histiobranchus bathybius* | Synaphobranchidae | - | - | [2]: two records from Bering Sea and Mexico; unclear if species is found in California | Excluded- cannot determine whether found in California | Not found in 1970s dataset (deep-sea fish) |
| Actinopterygii | *Thaumatichthys axeli* | Thaumatichthyidae | - | - | [2]: only two records of species | Excluded- too little information to determine range | Not found in 1970s dataset (deep-sea fish) |
| Chondrichthyes | *Alopias pelagicus* | Alopiidae | - | - | [1] give northern limit as "southern CA". Range within California is unclear | Yes | Not found in 1970s dataset |

| | | | | | | | |
|---|---|---|---|---|---|---|---|
| Chondrichthyes | *Carcharhinus limbatus* | Carcharhinidae | - | - | [2]: "unconfirmed records from California". Not mentioned in [1] | Excluded- cannot determine whether found in California | Not found in 1970s dataset |
| Chondrichthyes | *Carcharhinus longimanus* | Carcharhinidae | 32 | S of CA | [1] give northern limit as "perhaps Gaviota". If true, this would shift the range northward from 32 to 34 degrees. However, northern limit is uncertain. | yes | Unclear |
| Chondrichthyes | *Galeocerdo cuvier* | Carcharhinidae | 33 | S of CA | [1] claim this species ranges from Gulf of Alaska to Peru. However, [2] claim records in Alaska are unverified, give northern limit as "southern California". | Excluded- cannot determine whether range extends beyond California | Unclear |
| Chondrichthyes | *Hydrolagus trolli* | Chimaeridae | - | - | [1] claim it is normally a Central Pacific species, but it has been spotted in southern CA from ROV. These sightings are unconfirmed to be this species | Excluded- cannot determine whether found in California | Not found in 1970s dataset |
| Chondrichthyes | *Centroscyllium nigrum* | Etmopteridae | - | - | [1] gives northern limit as "southern California." Range within California is unclear | Yes | Not found in 1970s dataset |
| Chondrichthyes | *Isurus paucus* | Lamnidae | - | - | [1]: Only two records known from CA waters, range is unclear within California | Yes | Not found in 1970s dataset |
| Chondrichthyes | *Megachasma pelagios* | Megachasmidae | - | - | [1]: "occasional off CA". Range within California is unclear | Yes | Not found in 1970s dataset |
| Chondrichthyes | *Mitsukurina owstoni* | Mitsukurinidae | - | - | [1] cite one sighting in San Clemente, but otherwise not known from this region. Unclear if this is a vagrant species | Excluded- cannot determine whether found in California | Not found in 1970s dataset |
| Chondrichthyes | *Mobula japonica (Mobula mobular)* | Mobulidae | 34 | S of CA | [1] give northern limit as "central California". Cannot assign to latitudinal bands | yes | Unclear |
| Chondrichthyes | *Sphryna zygaena* | Sphyrnidae | 37 | S of CA | [1] give northern limit as "central California". Cannot assign to latitudinal bands | yes | Unclear |

| | | | | | | | |
|---|---|---|---|---|---|---|---|
| Chondrichthyes | *Sphyrna tiburo* | Sphyrnidae | 33 | S of CA | [1] claim it is historically found from San Diego to Peru. [2] claims it is formerly found in San Diego. Unclear if it is extirpated from San Diego (and thus no longer found in California waters) | Excluded- cannot determine whether still found in California | Unclear |

**Table S4:** Region-of-origin of California ray-finned fishes as inferred from BioGeoBEARS analyses.

| Species | Family | Order | habitat | # maps inferring northern origin | # maps inferring southern origin | # maps indet. | region of origin coded in plots | notes |
|---|---|---|---|---|---|---|---|---|
| *Acanthurus xanthopterus* | Acanthuridae | Acanthuriformes | demersal | 0 | 97 | 3 | S | |
| *Prognathodes falcifer* | Chaetodontidae | Acanthuriformes | demersal | 0 | 97 | 3 | S | |
| *Lobotes pacificus* | Lobotidae | Acanthuriformes | pelagic | 0 | 91 | 9 | S | |
| *Luvarus imperialis* | Luvaridae | Acanthuriformes | pelagic | 20 | 73 | 7 | S | |
| *Pomacanthus zonipectus* | Pomacanthidae | Acanthuriformes | demersal | 0 | 96 | 4 | S | |
| *Acipenser medirostris* | Acipenseridae | Acipenseriformes | diadromous | 36 | 47 | 17 | unclear | |
| *Acipenser transmontanus* | Acipenseridae | Acipenseriformes | diadromous | 36 | 32 | 32 | unclear | |
| *Pseudopentaceros wheeleri* | Pentacerotidae | Acropomatiformes | both demersal and pelagic | 93 | 0 | 7 | N | |
| *Stereolepis gigas* | Polyprionidae | Acropomatiformes | demersal | 0 | 97 | 3 | S | |
| *Albula vulpes* | Albulidae | Albuliformes | demersal | 0 | 93 | 7 | S | |
| *Muraena argus* | Muraenidae | Anguilliformes | demersal | 0 | 99 | 1 | S | |
| *Myrophis vafer* | Ophichthidae | Anguilliformes | demersal | 0 | 90 | 10 | S | |
| *Ophichthus triserialis* | Ophichthidae | Anguilliformes | demersal | 11 | 85 | 4 | S | |
| *Ophichthus zophochir* | Ophichthidae | Anguilliformes | demersal | 3 | 95 | 2 | S | |
| *Argentina sialis* | Argentinidae | Argentiniformes | demersal | 58 | 39 | 3 | unclear | |
| *Atherinops affinis* | Atherinopsidae | Atheriniformes | pelagic | 6 | 70 | 24 | S | |
| *Atherinopsis californiensis* | Atherinopsidae | Atheriniformes | pelagic | 6 | 70 | 24 | S | |
| *Leuresthes tenuis* | Atherinopsidae | Atheriniformes | pelagic | 6 | 71 | 23 | S | |
| *Synodus lucioceps* | Synodontidae | Aulopiformes | demersal | 37 | 58 | 5 | unclear | |
| *Porichthys myriaster* | Batrachoididae | Batrachoidiformes | demersal | 0 | 98 | 2 | S | |
| *Porichthys notatus* | Batrachoididae | Batrachoidiformes | demersal | 0 | 98 | 2 | S | |
| *Strongylura exilis* | Belonidae | Beloniformes | pelagic | 0 | 58 | 42 | probably S | |
| *Cheilopogon pinnatibarbatus californicus* | Exocoetidae | Beloniformes | pelagic | 36 | 63 | 1 | unclear | |
| *Fodiator acutus* | Exocoetidae | Beloniformes | pelagic | 0 | 96 | 4 | S | |
| *Euleptorhamphus viridis* | Hemiramphidae | Beloniformes | pelagic | 0 | 88 | 12 | S | |
| *Hemiramphus saltator* | Hemiramphidae | Beloniformes | pelagic | 0 | 98 | 2 | S | |
| *Cololabis saira* | Scomberesocidae | Beloniformes | pelagic | 47 | 28 | 25 | unclear | |
| *Hypsoblennius gentilis* | Blenniidae | Blenniiformes | demersal | 0 | 95 | 5 | S | |
| *Hypsoblennius gilberti* | Blenniidae | Blenniiformes | demersal | 0 | 97 | 3 | S | |

| | | | | | | | |
|---|---|---|---|---|---|---|---|
| *Hypsoblennius jenkinsi* | Blenniidae | Blenniiformes | demersal | 0 | 94 | 6 | S |
| *Ophioblennius steindachneri* | Blenniidae | Blenniiformes | demersal | 1 | 94 | 5 | S |
| *Plagiotremus azaleus* | Blenniidae | Blenniiformes | demersal | 0 | 97 | 3 | S |
| *Chaenopsis alepidota* | Chaenopsidae | Blenniiformes | demersal | 0 | 95 | 5 | S |
| *Neoclinus blanchardi* | Chaenopsidae | Blenniiformes | demersal | 0 | 99 | 1 | S |
| *Gibbonsia elegans* | Clinidae | Blenniiformes | demersal | 28 | 67 | 5 | unclear |
| *Gibbonsia metzi* | Clinidae | Blenniiformes | demersal | 28 | 67 | 5 | unclear |
| *Gibbonsia montereyensis* | Clinidae | Blenniiformes | demersal | 28 | 67 | 5 | unclear |
| *Heterostichus rostratus* | Clinidae | Blenniiformes | demersal | 28 | 67 | 5 | unclear |
| *Alloclinus holderi* | Labrisomidae | Blenniiformes | demersal | 0 | 94 | 6 | S |
| *Labrisomus xanti* | Labrisomidae | Blenniiformes | demersal | 0 | 100 | 0 | S |
| *Paraclinus integripinnis* | Labrisomidae | Blenniiformes | demersal | 0 | 97 | 3 | S |
| *Synchiropus atrilabiatus* | Callionymidae | Callionymiformes | demersal | 0 | 89 | 11 | S |
| *Polydactylus approximans* | Polynemidae | Carangaria/misc | demersal | 1 | 68 | 31 | probably S |
| *Sphyraena argentea* | Sphyraenidae | Carangaria/misc | pelagic | 40 | 52 | 8 | unclear |
| *Sphyraena ensis* | Sphyraenidae | Carangaria/misc | pelagic | 0 | 90 | 10 | S |
| *Caranx caballus* | Carangidae | Carangiformes | pelagic | 0 | 97 | 3 | S |
| *Caranx caninus* | Carangidae | Carangiformes | pelagic | 0 | 89 | 11 | S |
| *Caranx sexfasciatus* | Carangidae | Carangiformes | pelagic | 0 | 52 | 48 | probably S |
| *Caranx vinctus* | Carangidae | Carangiformes | pelagic | 0 | 89 | 11 | S |
| *Chloroscombrus orqueta* | Carangidae | Carangiformes | pelagic | 0 | 91 | 9 | S |
| *Decapterus muroadsi* | Carangidae | Carangiformes | pelagic | 0 | 100 | 0 | S |
| *Naucrates ductor* | Carangidae | Carangiformes | pelagic | 40 | 53 | 7 | unclear |
| *Selar crumenophthalmus* | Carangidae | Carangiformes | pelagic | 0 | 98 | 2 | S |
| *Selene brevoortii* | Carangidae | Carangiformes | demersal | 0 | 96 | 4 | S |
| *Selene peruviana* | Carangidae | Carangiformes | demersal | 0 | 95 | 5 | S |
| *Seriola lalandi* | Carangidae | Carangiformes | pelagic | 40 | 54 | 6 | unclear |
| *Seriola rivoliana* | Carangidae | Carangiformes | pelagic | 0 | 94 | 6 | S |
| *Trachinotus paitensis* | Carangidae | Carangiformes | pelagic | 1 | 91 | 8 | S |
| *Trachinotus rhodopus* | Carangidae | Carangiformes | demersal | 0 | 93 | 7 | S |
| *Trachurus symmetricus* | Carangidae | Carangiformes | pelagic | 48 | 47 | 5 | unclear |
| *Uraspis secunda* | Carangidae | Carangiformes | demersal | 0 | 97 | 3 | S |
| *Coryphaena equiselis* | Coryphaenidae | Carangiformes | pelagic | 0 | 98 | 2 | S |
| *Coryphaena hippurus* | Coryphaenidae | Carangiformes | pelagic | 0 | 98 | 2 | S |
| *Echeneis naucrates* | Echeneidae | Carangiformes | pelagic | 0 | 98 | 2 | S |
| *Phtheirichthys lineatus* | Echeneidae | Carangiformes | pelagic | 0 | 98 | 2 | S |
| *Remora albescens* | Echeneidae | Carangiformes | pelagic | 0 | 98 | 2 | S |
| *Remora australis* | Echeneidae | Carangiformes | pelagic | 0 | 98 | 2 | S |

| | | | | | | | |
|---|---|---|---|---|---|---|---|
| *Remora brachyptera* | Echeneidae | Carangiformes | pelagic | 0 | 98 | 2 | S |
| *Remora osteochir* | Echeneidae | Carangiformes | pelagic | 0 | 98 | 2 | S |
| *Remora remora* | Echeneidae | Carangiformes | pelagic | 0 | 98 | 2 | S |
| *Istiompax indica* | Istiophoridae | Carangiformes | pelagic | 4 | 91 | 5 | S |
| *Istiophorus platypterus* | Istiophoridae | Carangiformes | pelagic | 4 | 91 | 5 | S |
| *Kajikia audax* | Istiophoridae | Carangiformes | pelagic | 4 | 91 | 5 | S |
| *Makaira mazara* | Istiophoridae | Carangiformes | pelagic | 4 | 91 | 5 | S |
| *Tetrapturus angustirostris* | Istiophoridae | Carangiformes | pelagic | 4 | 91 | 5 | S |
| *Xiphias gladius* | Xiphiidae | Carangiformes | pelagic | 4 | 91 | 5 | S |
| *Girella nigricans* | Girellidae | Centrarchiformes | demersal | 46 | 53 | 1 | unclear |
| *Kyphosus analogus* | Kyphosidae | Centrarchiformes | demersal | 0 | 93 | 7 | S |
| *Kyphosus azurea* | Kyphosidae | Centrarchiformes | demersal | 0 | 94 | 6 | S |
| *Medialuna californiensis* | Scorpididae | Centrarchiformes | demersal | 40 | 54 | 6 | unclear |
| *Clupea harengus* | Clupeidae | Clupeiformes | pelagic | 51 | 37 | 12 | unclear |
| *Harengula thrissina* | Clupeidae | Clupeiformes | pelagic | 0 | 81 | 19 | S |
| *Opisthonema libertate* | Clupeidae | Clupeiformes | pelagic | 0 | 98 | 2 | S |
| *Opisthonema medirastre* | Clupeidae | Clupeiformes | pelagic | 0 | 98 | 2 | S |
| *Sardinops sagax* | Clupeidae | Clupeiformes | pelagic | 53 | 39 | 8 | unclear |
| *Etrumeus sadina* | Dussumieriidae | Clupeiformes | pelagic | 0 | 93 | 7 | S |
| *Anchoa compressa* | Engraulidae | Clupeiformes | pelagic | 0 | 88 | 12 | S |
| *Anchoa delicatissima* | Engraulidae | Clupeiformes | pelagic | 0 | 64 | 36 | probably S |
| *Engraulis mordax* | Engraulidae | Clupeiformes | pelagic | 47 | 46 | 7 | unclear |
| *Fundulus parvipinnis* | Fundulidae | Cyprinodontiformes | demersal | 0 | 52 | 48 | probably S |
| *Elops affinis* | Elopidae | Elopiformes | pelagic | 0 | 54 | 46 | probably S |
| *Eucinostomus currani* | Gerreidae | Eupercaria/misc | demersal | 0 | 63 | 37 | probably S |
| *Conodon serrifer* | Haemulidae | Eupercaria/misc | demersal | 0 | 99 | 1 | S |
| *Haemulon flaviguttatum* | Haemulidae | Eupercaria/misc | demersal | 0 | 100 | 0 | S |
| *Haemulopsis axillaris* | Haemulidae | Eupercaria/misc | demersal | 0 | 100 | 0 | S |
| *Haemulopsis elongatus* | Haemulidae | Eupercaria/misc | demersal | 0 | 100 | 0 | S |
| *Microlepidotus inornatus* | Haemulidae | Eupercaria/misc | demersal | 0 | 100 | 0 | S |
| *Xenistius californiensis* | Haemulidae | Eupercaria/misc | pelagic | 1 | 95 | 4 | S |
| *Decodon melasma* | Labridae | Eupercaria/misc | demersal | 1 | 90 | 9 | S |
| *Halichoeres semicinctus* | Labridae | Eupercaria/misc | demersal | 0 | 96 | 4 | S |
| *Nicholsina denticulata* | Labridae | Eupercaria/misc | demersal | 1 | 95 | 4 | S |
| *Oxyjulis californica* | Labridae | Eupercaria/misc | demersal | 0 | 96 | 4 | S |
| *Semicossyphus pulcher* | Labridae | Eupercaria/misc | demersal | 0 | 96 | 4 | S |
| *Caulolatilus affinis* | Latilidae | Eupercaria/misc | demersal | 3 | 92 | 5 | S |
| *Caulolatilus princeps* | Latilidae | Eupercaria/misc | demersal | 3 | 92 | 5 | S |

| | | | | | | | |
|---|---|---|---|---|---|---|---|
| *Lutjanus argentiventris* | Lutjanidae | Eupercaria/misc | demersal | 0 | 94 | 6 | S |
| *Lutjanus colorado* | Lutjanidae | Eupercaria/misc | demersal | 0 | 72 | 28 | S |
| *Lutjanus novemfasciatus* | Lutjanidae | Eupercaria/misc | demersal | 0 | 72 | 28 | S |
| *Lutjanus peru* | Lutjanidae | Eupercaria/misc | demersal | 0 | 97 | 3 | S |
| *Heteropriacanthus cruentatus* | Priacanthidae | Eupercaria/misc | demersal | 0 | 100 | 0 | S |
| *Pristigenys serrula* | Priacanthidae | Eupercaria/misc | demersal | 40 | 52 | 8 | unclear |
| *Atractoscion nobilis* | Sciaenidae | Eupercaria/misc | demersal | 44 | 52 | 4 | unclear |
| *Cheilotrema saturnum* | Sciaenidae | Eupercaria/misc | demersal | 0 | 96 | 4 | S |
| *Cynoscion parvipinnis* | Sciaenidae | Eupercaria/misc | demersal | 0 | 91 | 9 | S |
| *Genyonemus lineatus* | Sciaenidae | Eupercaria/misc | demersal | 9 | 82 | 9 | S |
| *Menticirrhus undulatus* | Sciaenidae | Eupercaria/misc | demersal | 0 | 88 | 12 | S |
| *Roncador stearnsii* | Sciaenidae | Eupercaria/misc | demersal | 9 | 82 | 9 | S |
| *Seriphus politus* | Sciaenidae | Eupercaria/misc | demersal | 9 | 82 | 9 | S |
| *Umbrina roncador* | Sciaenidae | Eupercaria/misc | demersal | 0 | 95 | 5 | S |
| *Calamus brachysomus* | Sparidae | Eupercaria/misc | demersal | 0 | 96 | 4 | S |
| *Gadus macrocephalus* | Gadidae | Gadiformes | both demersal and pelagic | 99 | 0 | 1 | N |
| *Microgadus proximus* | Gadidae | Gadiformes | demersal | 98 | 0 | 2 | N |
| *Theragra chalcogramma* | Gadidae | Gadiformes | both demersal and pelagic | 99 | 0 | 1 | N |
| *Coryphaenoides acrolepis* | Macrouridae | Gadiformes | both demersal and pelagic | 97 | 0 | 3 | N |
| *Merluccius productus* | Merlucciidae | Gadiformes | pelagic | 43 | 55 | 2 | unclear |
| *Antimora microlepis* | Moridae | Gadiformes | demersal | 72 | 20 | 8 | N |
| *Physiculus rastrelliger* | Moridae | Gadiformes | demersal | 46 | 48 | 6 | unclear |
| *Gobiesox maeandricus* | Gobiesocidae | Gobiesociformes | demersal | 0 | 100 | 0 | S |
| *Gobiesox rhessodon* | Gobiesocidae | Gobiesociformes | demersal | 0 | 100 | 0 | S |
| *Dormitator latifrons* | Eleotridae | Gobiiformes | demersal | 0 | 73 | 27 | S |
| *Clevelandia ios* | Gobiidae | Gobiiformes | demersal | 0 | 89 | 11 | S |
| *Ctenogobius sagittula* | Gobiidae | Gobiiformes | demersal | 0 | 79 | 21 | S |
| *Eucyclogobius newberryi* | Gobiidae | Gobiiformes | demersal | 0 | 92 | 8 | S |
| *Gillichthys mirabilis* | Gobiidae | Gobiiformes | demersal | 0 | 93 | 7 | S |
| *Lepidogobius lepidus* | Gobiidae | Gobiiformes | demersal | 0 | 92 | 8 | S |
| *Lethops connectens* | Gobiidae | Gobiiformes | demersal | 0 | 89 | 11 | S |
| *Lythrypnus dalli* | Gobiidae | Gobiiformes | demersal | 0 | 94 | 6 | S |
| *Lythrypnus zebra* | Gobiidae | Gobiiformes | demersal | 0 | 97 | 3 | S |
| *Quietula y-cauda* | Gobiidae | Gobiiformes | demersal | 0 | 89 | 11 | S |
| *Rhinogobiops nicholsii* | Gobiidae | Gobiiformes | demersal | 48 | 44 | 8 | unclear |

| | | | | | | | |
|---|---|---|---|---|---|---|---|
| *Typhlogobius californiensis* | Gobiidae | Gobiiformes | demersal | 0 | 92 | 8 | S |
| *Chanos chanos* | Chanidae | Gonorynchiformes | pelagic | 3 | 79 | 18 | S |
| *Apogon guadalupensis* | Apogonidae | Kurtiformes | demersal | 0 | 100 | 0 | S |
| *Apogon pacificus* | Apogonidae | Kurtiformes | demersal | 0 | 100 | 0 | S |
| *Lophotus capellei* | Lophotidae | Lampriformes | pelagic | 18 | 15 | 67 | unclear |
| *Trachipterus altivelis* | Trachipteridae | Lampriformes | pelagic | 47 | 47 | 6 | unclear |
| *Zu cristatus* | Trachipteridae | Lampriformes | pelagic | 1 | 93 | 6 | S |
| *Fowlerichthys avalonis* | Antennariidae | Lophiiformes | demersal | 0 | 94 | 6 | S |
| *Lophiodes caulinaris* | Lophiidae | Lophiiformes | demersal | 0 | 96 | 4 | S |
| *Lophiodes spilurus* | Lophiidae | Lophiiformes | demersal | 0 | 96 | 4 | S |
| *Zalieutes elater* | Ogcocephalidae | Lophiiformes | demersal | 49 | 41 | 10 | unclear |
| *Mugil cephalus* | Mugilidae | Mugiliformes | diadromous | 0 | 67 | 33 | probably S |
| *Mugil curema* | Mugilidae | Mugiliformes | pelagic | 0 | 61 | 39 | probably S |
| *Pseudupeneus grandisquamis* | Mullidae | Mulliformes | demersal | 0 | 98 | 2 | S |
| *Brosmophycis marginata* | Bythitidae | Ophidiiformes | demersal | 49 | 46 | 5 | unclear |
| *Chilara taylori* | Ophidiidae | Ophidiiformes | demersal | 53 | 41 | 6 | unclear |
| *Allosmerus elongatus* | Osmeridae | Osmeriformes | demersal | 87 | 0 | 13 | N |
| *Hypomesus pretiosus* | Osmeridae | Osmeriformes | demersal | 68 | 0 | 32 | probably N |
| *Spirinchus starksi* | Osmeridae | Osmeriformes | demersal | 87 | 0 | 13 | N |
| *Spirinchus thaleichthys* | Osmeridae | Osmeriformes | diadromous | 87 | 0 | 13 | N |
| *Thaleichthys pacificus* | Osmeridae | Osmeriformes | diadromous | 87 | 0 | 13 | N |
| *Amphistichus koelzi* | Embiotocidae | Ovalentaria/misc | demersal | 8 | 82 | 10 | S |
| *Amphistichus rhodoterus* | Embiotocidae | Ovalentaria/misc | demersal | 8 | 82 | 10 | S |
| *Brachyistius frenatus* | Embiotocidae | Ovalentaria/misc | demersal | 8 | 82 | 10 | S |
| *Cymatogaster aggregata* | Embiotocidae | Ovalentaria/misc | demersal | 8 | 82 | 10 | S |
| *Embiotoca jacksoni* | Embiotocidae | Ovalentaria/misc | demersal | 8 | 82 | 10 | S |
| *Embiotoca lateralis* | Embiotocidae | Ovalentaria/misc | demersal | 8 | 82 | 10 | S |
| *Hyperprosopon anale* | Embiotocidae | Ovalentaria/misc | pelagic | 8 | 82 | 10 | S |
| *Hyperprosopon argenteum* | Embiotocidae | Ovalentaria/misc | demersal | 8 | 82 | 10 | S |
| *Hyperprosopon ellipticum* | Embiotocidae | Ovalentaria/misc | demersal | 8 | 82 | 10 | S |
| *Hypsurus caryi* | Embiotocidae | Ovalentaria/misc | demersal | 8 | 82 | 10 | S |
| *Micrometrus minimus* | Embiotocidae | Ovalentaria/misc | demersal | 8 | 82 | 10 | S |
| *Phanerodon atripes* | Embiotocidae | Ovalentaria/misc | demersal | 8 | 82 | 10 | S |
| *Phanerodon furcatus* | Embiotocidae | Ovalentaria/misc | demersal | 8 | 82 | 10 | S |
| *Rhacochilus toxotes* | Embiotocidae | Ovalentaria/misc | demersal | 8 | 82 | 10 | S |
| *Rhacochilus vacca* | Embiotocidae | Ovalentaria/misc | demersal | 8 | 82 | 10 | S |
| *Zalembius rosaceus* | Embiotocidae | Ovalentaria/misc | demersal | 8 | 82 | 10 | S |
| *Abudefduf troschelii* | Pomacentridae | Ovalentaria/misc | demersal | 0 | 93 | 7 | S |

| | | | | | | | |
|---|---|---|---|---|---|---|---|
| *Azurina hirundo* | Pomacentridae | Ovalentaria/misc | demersal | 0 | 100 | 0 | S |
| *Chromis alta* | Pomacentridae | Ovalentaria/misc | demersal | 0 | 95 | 5 | S |
| *Chromis punctipinnis* | Pomacentridae | Ovalentaria/misc | demersal | 0 | 96 | 4 | S |
| *Hypsypops rubicundus* | Pomacentridae | Ovalentaria/misc | demersal | 0 | 99 | 1 | S |
| *Agonopsis sterletus* | Agonidae | Perciformes/Cottoidei | demersal | 98 | 0 | 2 | N |
| *Agonopsis vulsa* | Agonidae | Perciformes/Cottoidei | demersal | 98 | 0 | 2 | N |
| *Anoplagonus inermis* | Agonidae | Perciformes/Cottoidei | demersal | 100 | 0 | 0 | N |
| *Bathyagonus alascanus* | Agonidae | Perciformes/Cottoidei | demersal | 98 | 0 | 2 | N |
| *Bathyagonus infraspinatus* | Agonidae | Perciformes/Cottoidei | demersal | 98 | 0 | 2 | N |
| *Bathyagonus nigripinnis* | Agonidae | Perciformes/Cottoidei | demersal | 98 | 0 | 2 | N |
| *Bathyagonus pentacanthus* | Agonidae | Perciformes/Cottoidei | demersal | 98 | 0 | 2 | N |
| *Blepsias cirrhosus* | Agonidae | Perciformes/Cottoidei | demersal | 100 | 0 | 0 | N |
| *Nautichthys oculofasciatus* | Agonidae | Perciformes/Cottoidei | demersal | 96 | 1 | 3 | N |
| *Odontopyxis trispinosa* | Agonidae | Perciformes/Cottoidei | demersal | 98 | 0 | 2 | N |
| *Podothecus accipenserinus* | Agonidae | Perciformes/Cottoidei | demersal | 100 | 0 | 0 | N |
| *Stellerina xyosterna* | Agonidae | Perciformes/Cottoidei | demersal | 98 | 0 | 2 | N |
| *Xeneretmus latifrons* | Agonidae | Perciformes/Cottoidei | demersal | 98 | 0 | 2 | N |
| *Xeneretmus leiops* | Agonidae | Perciformes/Cottoidei | demersal | 98 | 0 | 2 | N |
| *Anoplopoma fimbria* | Anoplopomatidae | Perciformes/Cottoidei | demersal | 95 | 2 | 3 | N |
| *Erilepis zonifer* | Anoplopomatidae | Perciformes/Cottoidei | demersal | 95 | 2 | 3 | N |
| *Artedius corallinus* | Cottidae | Perciformes/Cottoidei | demersal | 100 | 0 | 0 | N |
| *Artedius fenestralis* | Cottidae | Perciformes/Cottoidei | demersal | 100 | 0 | 0 | N |
| *Artedius harringtoni* | Cottidae | Perciformes/Cottoidei | demersal | 100 | 0 | 0 | N |
| *Artedius lateralis* | Cottidae | Perciformes/Cottoidei | demersal | 100 | 0 | 0 | N |
| *Artedius notospilotus* | Cottidae | Perciformes/Cottoidei | demersal | 100 | 0 | 0 | N |
| *Chitonotus pugetensis* | Cottidae | Perciformes/Cottoidei | demersal | 100 | 0 | 0 | N |
| *Clinocottus acuticeps* | Cottidae | Perciformes/Cottoidei | demersal | 100 | 0 | 0 | N |
| *Clinocottus analis* | Cottidae | Perciformes/Cottoidei | demersal | 100 | 0 | 0 | N |
| *Clinocottus embryum* | Cottidae | Perciformes/Cottoidei | demersal | 100 | 0 | 0 | N |
| *Clinocottus globiceps* | Cottidae | Perciformes/Cottoidei | demersal | 100 | 0 | 0 | N |
| *Clinocottus recalvus* | Cottidae | Perciformes/Cottoidei | demersal | 100 | 0 | 0 | N |
| *Cottus asper* | Cottidae | Perciformes/Cottoidei | diadromous | 57 | 0 | 43 | probably N |
| *Enophrys bison* | Cottidae | Perciformes/Cottoidei | demersal | 79 | 0 | 21 | N |
| *Enophrys taurina* | Cottidae | Perciformes/Cottoidei | demersal | 79 | 0 | 21 | N |
| *Hemilepidotus hemilepidotus* | Cottidae | Perciformes/Cottoidei | demersal | 99 | 0 | 1 | N |
| *Icelinus burchami* | Cottidae | Perciformes/Cottoidei | demersal | 100 | 0 | 0 | N |
| *Icelinus cavifrons* | Cottidae | Perciformes/Cottoidei | demersal | 99 | 1 | 0 | N |
| *Icelinus filamentosus* | Cottidae | Perciformes/Cottoidei | demersal | 99 | 1 | 0 | N |

| | | | | | | | |
|---|---|---|---|---|---|---|---|
| *Icelinus fimbriatus* | Cottidae | Perciformes/Cottoidei | demersal | 100 | 0 | 0 | N |
| *Icelinus oculatus* | Cottidae | Perciformes/Cottoidei | demersal | 100 | 0 | 0 | N |
| *Icelinus quadriseriatus* | Cottidae | Perciformes/Cottoidei | demersal | 99 | 1 | 0 | N |
| *Icelinus tenuis* | Cottidae | Perciformes/Cottoidei | demersal | 99 | 1 | 0 | N |
| *Jordania zonope* | Cottidae | Perciformes/Cottoidei | demersal | 97 | 1 | 2 | N |
| *Leiocottus hirundo* | Cottidae | Perciformes/Cottoidei | demersal | 100 | 0 | 0 | N |
| *Leptocottus armatus* | Cottidae | Perciformes/Cottoidei | demersal | 69 | 10 | 21 | probably N |
| *Oligocottus maculosus* | Cottidae | Perciformes/Cottoidei | demersal | 100 | 0 | 0 | N |
| *Oligocottus rimensis* | Cottidae | Perciformes/Cottoidei | demersal | 100 | 0 | 0 | N |
| *Oligocottus rubellio* | Cottidae | Perciformes/Cottoidei | demersal | 100 | 0 | 0 | N |
| *Oligocottus snyderi* | Cottidae | Perciformes/Cottoidei | demersal | 100 | 0 | 0 | N |
| *Orthonopias triacis* | Cottidae | Perciformes/Cottoidei | demersal | 100 | 0 | 0 | N |
| *Radulinus asprellus* | Cottidae | Perciformes/Cottoidei | demersal | 100 | 0 | 0 | N |
| *Radulinus boleoides* | Cottidae | Perciformes/Cottoidei | demersal | 100 | 0 | 0 | N |
| *Ruscarius creaseri* | Cottidae | Perciformes/Cottoidei | demersal | 100 | 0 | 0 | N |
| *Scorpaenichthys marmoratus* | Cottidae | Perciformes/Cottoidei | demersal | 97 | 1 | 2 | N |
| *Zesticelus profundorum* | Cottidae | Perciformes/Cottoidei | demersal | 98 | 2 | 0 | N |
| *Hexagrammos decagrammus* | Hexagrammidae | Perciformes/Cottoidei | demersal | 95 | 2 | 3 | N |
| *Hexagrammos lagocephalus* | Hexagrammidae | Perciformes/Cottoidei | demersal | 95 | 2 | 3 | N |
| *Hexagrammos stelleri* | Hexagrammidae | Perciformes/Cottoidei | demersal | 95 | 2 | 3 | N |
| *Ophiodon elongatus* | Hexagrammidae | Perciformes/Cottoidei | demersal | 95 | 2 | 3 | N |
| *Pleurogrammus monopterygius* | Hexagrammidae | Perciformes/Cottoidei | demersal | 95 | 2 | 3 | N |
| *Careproctus melanurus* | Liparidae | Perciformes/Cottoidei | demersal | 96 | 1 | 3 | N |
| *Liparis florae* | Liparidae | Perciformes/Cottoidei | demersal | 96 | 1 | 3 | N |
| *Liparis fucensis* | Liparidae | Perciformes/Cottoidei | demersal | 98 | 0 | 2 | N |
| *Liparis mucosus* | Liparidae | Perciformes/Cottoidei | demersal | 96 | 1 | 3 | N |
| *Liparis pulchellus* | Liparidae | Perciformes/Cottoidei | demersal | 96 | 1 | 3 | N |
| *Liparis rutteri* | Liparidae | Perciformes/Cottoidei | demersal | 96 | 1 | 3 | N |
| *Lipariscus nanus* | Liparidae | Perciformes/Cottoidei | demersal | 96 | 1 | 3 | N |
| *Rhamphocottus richardsonii* | Rhamphocottidae | Perciformes/Cottoidei | demersal | 97 | 1 | 2 | N |
| *Oxylebius pictus* | Zaniolepididae | Perciformes/Cottoidei | demersal | 95 | 2 | 3 | N |
| *Zaniolepis frenata* | Zaniolepididae | Perciformes/Cottoidei | demersal | 95 | 2 | 3 | N |
| *Zaniolepis latipinnis* | Zaniolepididae | Perciformes/Cottoidei | demersal | 95 | 2 | 3 | N |
| *Aulorhynchus flavidus* | Aulorhynchidae | Perciformes/Gasterosteoidei | pelagic | 85 | 7 | 8 | N |
| *Scorpaena guttata* | Scorpaenidae | Perciformes/Scorpaenoidei | demersal | 0 | 93 | 7 | S |
| *Scorpaena mystes* | Scorpaenidae | Perciformes/Scorpaenoidei | demersal | 0 | 96 | 4 | S |
| *Scorpaenodes xyris* | Scorpaenidae | Perciformes/Scorpaenoidei | demersal | 0 | 93 | 7 | S |

| | | | | | | | |
|---|---|---|---|---|---|---|---|
| *Sebastes alutus* | Sebastidae | Perciformes/Scorpaenoidei | demersal | 96 | 0 | 4 | N |
| *Sebastes atrovirens* | Sebastidae | Perciformes/Scorpaenoidei | demersal | 96 | 0 | 4 | N |
| *Sebastes auriculatus* | Sebastidae | Perciformes/Scorpaenoidei | demersal | 96 | 0 | 4 | N |
| *Sebastes aurora* | Sebastidae | Perciformes/Scorpaenoidei | demersal | 96 | 0 | 4 | N |
| *Sebastes babcocki* | Sebastidae | Perciformes/Scorpaenoidei | demersal | 96 | 0 | 4 | N |
| *Sebastes borealis* | Sebastidae | Perciformes/Scorpaenoidei | demersal | 96 | 0 | 4 | N |
| *Sebastes brevispinis* | Sebastidae | Perciformes/Scorpaenoidei | demersal | 96 | 0 | 4 | N |
| *Sebastes carnatus* | Sebastidae | Perciformes/Scorpaenoidei | demersal | 96 | 0 | 4 | N |
| *Sebastes caurinus* | Sebastidae | Perciformes/Scorpaenoidei | demersal | 96 | 0 | 4 | N |
| *Sebastes chlorostictus* | Sebastidae | Perciformes/Scorpaenoidei | demersal | 96 | 0 | 4 | N |
| *Sebastes chrysomelas* | Sebastidae | Perciformes/Scorpaenoidei | demersal | 96 | 0 | 4 | N |
| *Sebastes constellatus* | Sebastidae | Perciformes/Scorpaenoidei | demersal | 96 | 0 | 4 | N |
| *Sebastes crameri* | Sebastidae | Perciformes/Scorpaenoidei | demersal | 100 | 0 | 0 | N |
| *Sebastes dallii* | Sebastidae | Perciformes/Scorpaenoidei | demersal | 96 | 0 | 4 | N |
| *Sebastes diploproa* | Sebastidae | Perciformes/Scorpaenoidei | demersal | 96 | 0 | 4 | N |
| *Sebastes elongatus* | Sebastidae | Perciformes/Scorpaenoidei | demersal | 96 | 0 | 4 | N |
| *Sebastes emphaeus* | Sebastidae | Perciformes/Scorpaenoidei | demersal | 96 | 0 | 4 | N |
| *Sebastes ensifer* | Sebastidae | Perciformes/Scorpaenoidei | demersal | 96 | 0 | 4 | N |
| *Sebastes entomelas* | Sebastidae | Perciformes/Scorpaenoidei | demersal | 96 | 0 | 4 | N |
| *Sebastes eos* | Sebastidae | Perciformes/Scorpaenoidei | demersal | 96 | 0 | 4 | N |
| *Sebastes flavidus* | Sebastidae | Perciformes/Scorpaenoidei | pelagic | 96 | 0 | 4 | N |
| *Sebastes gilli* | Sebastidae | Perciformes/Scorpaenoidei | demersal | 96 | 0 | 4 | N |
| *Sebastes helvomaculatus* | Sebastidae | Perciformes/Scorpaenoidei | demersal | 96 | 0 | 4 | N |
| *Sebastes hopkinsi* | Sebastidae | Perciformes/Scorpaenoidei | demersal | 96 | 0 | 4 | N |
| *Sebastes jordani* | Sebastidae | Perciformes/Scorpaenoidei | demersal | 96 | 0 | 4 | N |
| *Sebastes lentiginosus* | Sebastidae | Perciformes/Scorpaenoidei | demersal | 96 | 0 | 4 | N |
| *Sebastes macdonaldi* | Sebastidae | Perciformes/Scorpaenoidei | demersal | 96 | 0 | 4 | N |
| *Sebastes maliger* | Sebastidae | Perciformes/Scorpaenoidei | demersal | 96 | 0 | 4 | N |
| *Sebastes melanops* | Sebastidae | Perciformes/Scorpaenoidei | demersal | 96 | 0 | 4 | N |
| *Sebastes melanosema* | Sebastidae | Perciformes/Scorpaenoidei | demersal | 95 | 1 | 4 | N |
| *Sebastes melanostictus* | Sebastidae | Perciformes/Scorpaenoidei | demersal | 96 | 0 | 4 | N |
| *Sebastes melanostomus* | Sebastidae | Perciformes/Scorpaenoidei | demersal | 96 | 0 | 4 | N |
| *Sebastes miniatus* | Sebastidae | Perciformes/Scorpaenoidei | demersal | 96 | 0 | 4 | N |
| *Sebastes moseri* | Sebastidae | Perciformes/Scorpaenoidei | demersal | 96 | 0 | 4 | N |
| *Sebastes mystinus* | Sebastidae | Perciformes/Scorpaenoidei | pelagic | 96 | 0 | 4 | N |
| *Sebastes nebulosus* | Sebastidae | Perciformes/Scorpaenoidei | demersal | 96 | 0 | 4 | N |
| *Sebastes nigrocinctus* | Sebastidae | Perciformes/Scorpaenoidei | demersal | 96 | 0 | 4 | N |
| *Sebastes ovalis* | Sebastidae | Perciformes/Scorpaenoidei | demersal | 96 | 0 | 4 | N |

| | | | | | | | |
|---|---|---|---|---|---|---|---|
| *Sebastes paucispinis* | Sebastidae | Perciformes/Scorpaenoidei | demersal | 96 | 0 | 4 | N |
| *Sebastes phillipsi* | Sebastidae | Perciformes/Scorpaenoidei | demersal | 96 | 0 | 4 | N |
| *Sebastes pinniger* | Sebastidae | Perciformes/Scorpaenoidei | demersal | 96 | 0 | 4 | N |
| *Sebastes proriger* | Sebastidae | Perciformes/Scorpaenoidei | demersal | 96 | 0 | 4 | N |
| *Sebastes rastrelliger* | Sebastidae | Perciformes/Scorpaenoidei | demersal | 96 | 0 | 4 | N |
| *Sebastes reedi* | Sebastidae | Perciformes/Scorpaenoidei | demersal | 100 | 0 | 0 | N |
| *Sebastes rosenblatti* | Sebastidae | Perciformes/Scorpaenoidei | demersal | 96 | 0 | 4 | N |
| *Sebastes ruberrimus* | Sebastidae | Perciformes/Scorpaenoidei | demersal | 96 | 0 | 4 | N |
| *Sebastes rubrivinctus* | Sebastidae | Perciformes/Scorpaenoidei | demersal | 96 | 0 | 4 | N |
| *Sebastes rufinanus* | Sebastidae | Perciformes/Scorpaenoidei | demersal | 96 | 0 | 4 | N |
| *Sebastes rufus* | Sebastidae | Perciformes/Scorpaenoidei | demersal | 96 | 0 | 4 | N |
| *Sebastes saxicola* | Sebastidae | Perciformes/Scorpaenoidei | demersal | 96 | 0 | 4 | N |
| *Sebastes semicinctus* | Sebastidae | Perciformes/Scorpaenoidei | demersal | 96 | 0 | 4 | N |
| *Sebastes serranoides* | Sebastidae | Perciformes/Scorpaenoidei | demersal | 96 | 0 | 4 | N |
| *Sebastes serriceps* | Sebastidae | Perciformes/Scorpaenoidei | demersal | 96 | 0 | 4 | N |
| *Sebastes simulator* | Sebastidae | Perciformes/Scorpaenoidei | demersal | 96 | 0 | 4 | N |
| *Sebastes umbrosus* | Sebastidae | Perciformes/Scorpaenoidei | demersal | 96 | 0 | 4 | N |
| *Sebastes wilsoni* | Sebastidae | Perciformes/Scorpaenoidei | demersal | 96 | 0 | 4 | N |
| *Sebastes zacentrus* | Sebastidae | Perciformes/Scorpaenoidei | demersal | 96 | 0 | 4 | N |
| *Sebastolobus alascanus* | Sebastidae | Perciformes/Scorpaenoidei | demersal | 100 | 0 | 0 | N |
| *Sebastolobus altivelis* | Sebastidae | Perciformes/Scorpaenoidei | demersal | 100 | 0 | 0 | N |
| *Bellator xenisma* | Triglidae | Perciformes/Scorpaenoidei | demersal | 0 | 97 | 3 | S |
| *Prionotus stephanophrys* | Triglidae | Perciformes/Scorpaenoidei | demersal | 37 | 56 | 7 | unclear |
| *Dermatolepis dermatolepis* | Serranidae | Perciformes/Serranoidei | demersal | 0 | 97 | 3 | S |
| *Diplectrum maximum* | Serranidae | Perciformes/Serranoidei | demersal | 0 | 99 | 1 | S |
| *Epinephelus analogus* | Serranidae | Perciformes/Serranoidei | demersal | 0 | 94 | 6 | S |
| *Epinephelus labriformis* | Serranidae | Perciformes/Serranoidei | demersal | 0 | 96 | 4 | S |
| *Hemanthias signifer* | Serranidae | Perciformes/Serranoidei | demersal | 0 | 90 | 10 | S |
| *Hyporthodus acanthistius* | Serranidae | Perciformes/Serranoidei | demersal | 0 | 98 | 2 | S |
| *Hyporthodus niphobles* | Serranidae | Perciformes/Serranoidei | demersal | 0 | 98 | 2 | S |
| *Mycteroperca jordani* | Serranidae | Perciformes/Serranoidei | demersal | 0 | 100 | 0 | S |
| *Mycteroperca xenarcha* | Serranidae | Perciformes/Serranoidei | demersal | 0 | 97 | 3 | S |
| *Paralabrax auroguttatus* | Serranidae | Perciformes/Serranoidei | demersal | 0 | 96 | 4 | S |
| *Paralabrax clathratus* | Serranidae | Perciformes/Serranoidei | demersal | 0 | 96 | 4 | S |
| *Paralabrax maculatofasciatus* | Serranidae | Perciformes/Serranoidei | demersal | 0 | 96 | 4 | S |
| *Paralabrax nebulifer* | Serranidae | Perciformes/Serranoidei | demersal | 0 | 96 | 4 | S |
| *Paranthias colonus* | Serranidae | Perciformes/Serranoidei | pelagic | 1 | 94 | 5 | S |

| | | | | | | | |
|---|---|---|---|---|---|---|---|
| *Pronotogrammus multifasciatus* | Serranidae | Perciformes/Serranoidei | demersal | 0 | 97 | 3 | S |
| *Ammodytes personatus* | Ammodytidae | Perciformes/Uranoscopoidei | demersal | 96 | 0 | 4 | N |
| *Trichodon trichodon* | Trichodontidae | Perciformes/Uranoscopoidei | demersal | 96 | 1 | 3 | N |
| *Kathetostoma averruncus* | Uranoscopidae | Perciformes/Uranoscopoidei | demersal | 0 | 95 | 5 | S |
| *Anarrhichthys ocellatus* | Anarhichadidae | Perciformes/Zoarcoidei | demersal | 97 | 3 | 0 | N |
| *Rathbunella hypoplecta* | Bathymasteridae | Perciformes/Zoarcoidei | demersal | 95 | 2 | 3 | N |
| *Ronquilus jordani* | Bathymasteridae | Perciformes/Zoarcoidei | demersal | 95 | 2 | 3 | N |
| *Cebidichthys violaceus* | Cebidichthyidae | Perciformes/Zoarcoidei | demersal | 95 | 2 | 3 | N |
| *Cryptacanthodes aleutensis* | Cryptacanthodidae | Perciformes/Zoarcoidei | demersal | 95 | 2 | 3 | N |
| *Cryptacanthodes giganteus* | Cryptacanthodidae | Perciformes/Zoarcoidei | demersal | 95 | 2 | 3 | N |
| *Lumpenus sagitta* | Lumpenidae | Perciformes/Zoarcoidei | demersal | 100 | 0 | 0 | N |
| *Poroclinus rothrocki* | Lumpenidae | Perciformes/Zoarcoidei | demersal | 95 | 2 | 3 | N |
| *Kasatkia seigeli* | Opisthocentridae | Perciformes/Zoarcoidei | demersal | 33 | 2 | 65 | probably N |
| *Apodichthys flavidus* | Pholidae | Perciformes/Zoarcoidei | demersal | 95 | 2 | 3 | N |
| *Apodichthys fucorum* | Pholidae | Perciformes/Zoarcoidei | demersal | 95 | 2 | 3 | N |
| *Pholis clemensi* | Pholidae | Perciformes/Zoarcoidei | demersal | 95 | 2 | 3 | N |
| *Pholis laeta* | Pholidae | Perciformes/Zoarcoidei | demersal | 95 | 2 | 3 | N |
| *Pholis ornata* | Pholidae | Perciformes/Zoarcoidei | demersal | 95 | 2 | 3 | N |
| *Anoplarchus insignis* | Stichaeidae | Perciformes/Zoarcoidei | demersal | 95 | 2 | 3 | N |
| *Anoplarchus purpurescens* | Stichaeidae | Perciformes/Zoarcoidei | demersal | 95 | 2 | 3 | N |
| *Chirolophis decoratus* | Stichaeidae | Perciformes/Zoarcoidei | demersal | 97 | 0 | 3 | N |
| *Chirolophis nugator* | Stichaeidae | Perciformes/Zoarcoidei | demersal | 97 | 0 | 3 | N |
| *Esselenichthys carli* | Stichaeidae | Perciformes/Zoarcoidei | demersal | 95 | 2 | 3 | N |
| *Xiphister mucosus* | Stichaeidae | Perciformes/Zoarcoidei | demersal | 95 | 2 | 3 | N |
| *Zaprora silenus* | Zaproridae | Perciformes/Zoarcoidei | demersal | 95 | 2 | 3 | N |
| *Bothrocara brunneum* | Zoarcidae | Perciformes/Zoarcoidei | demersal | 73 | 0 | 27 | N |
| *Bothrocara molle* | Zoarcidae | Perciformes/Zoarcoidei | demersal | 73 | 0 | 27 | N |
| *Eucryphycus californicus* | Zoarcidae | Perciformes/Zoarcoidei | demersal | 73 | 0 | 27 | N |
| *Lycenchelys crotalinus* | Zoarcidae | Perciformes/Zoarcoidei | demersal | 73 | 0 | 27 | N |
| *Lycodapus fierasfer* | Zoarcidae | Perciformes/Zoarcoidei | both demersal and pelagic | 73 | 0 | 27 | N |
| *Lycodapus mandibularis* | Zoarcidae | Perciformes/Zoarcoidei | pelagic | 74 | 0 | 26 | N |
| *Lycodes brevipes* | Zoarcidae | Perciformes/Zoarcoidei | demersal | 97 | 0 | 3 | N |
| *Lycodes cortezianus* | Zoarcidae | Perciformes/Zoarcoidei | demersal | 73 | 0 | 27 | N |
| *Lycodes diapterus* | Zoarcidae | Perciformes/Zoarcoidei | demersal | 75 | 0 | 25 | N |

| | | | | | | | |
|---|---|---|---|---|---|---|---|
| *Lycodes pacificus* | Zoarcidae | Perciformes/Zoarcoidei | demersal | 73 | 0 | 27 | N |
| *Melanostigma pammelas* | Zoarcidae | Perciformes/Zoarcoidei | pelagic | 45 | 51 | 4 | unclear |
| *Citharichthys sordidus* | Cyclopsettidae | Pleuronectiformes | demersal | 0 | 100 | 0 | S |
| *Citharichthys stigmaeus* | Cyclopsettidae | Pleuronectiformes | demersal | 1 | 99 | 0 | S |
| *Citharichthys xanthostigma* | Cyclopsettidae | Pleuronectiformes | demersal | 0 | 100 | 0 | S |
| *Symphurus atricaudus* | Cynoglossidae | Pleuronectiformes | demersal | 46 | 49 | 5 | unclear |
| *Hippoglossina stomata* | Paralichthyidae | Pleuronectiformes | demersal | 1 | 96 | 3 | S |
| *Paralichthys californicus* | Paralichthyidae | Pleuronectiformes | demersal | 94 | 0 | 6 | N |
| *Xystreurys liolepis* | Paralichthyidae | Pleuronectiformes | demersal | 0 | 96 | 4 | S |
| *Atheresthes stomias* | Pleuronectidae | Pleuronectiformes | demersal | 55 | 41 | 4 | unclear |
| *Eopsetta jordani* | Pleuronectidae | Pleuronectiformes | demersal | 50 | 46 | 4 | unclear |
| *Glyptocephalus zachirus* | Pleuronectidae | Pleuronectiformes | demersal | 50 | 46 | 4 | unclear |
| *Hippoglossoides elassodon* | Pleuronectidae | Pleuronectiformes | demersal | 97 | 0 | 3 | N |
| *Hippoglossus stenolepis* | Pleuronectidae | Pleuronectiformes | demersal | 50 | 46 | 4 | unclear |
| *Hypsopsetta guttulata* | Pleuronectidae | Pleuronectiformes | demersal | 50 | 46 | 4 | unclear |
| *Isopsetta isolepis* | Pleuronectidae | Pleuronectiformes | demersal | 93 | 7 | 0 | N |
| *Lepidopsetta bilineata* | Pleuronectidae | Pleuronectiformes | demersal | 93 | 7 | 0 | N |
| *Lyopsetta exilis* | Pleuronectidae | Pleuronectiformes | demersal | 50 | 46 | 4 | unclear |
| *Microstomus pacificus* | Pleuronectidae | Pleuronectiformes | demersal | 50 | 46 | 4 | unclear |
| *Parophrys vetulus* | Pleuronectidae | Pleuronectiformes | demersal | 93 | 7 | 0 | N |
| *Platichthys stellatus* | Pleuronectidae | Pleuronectiformes | demersal | 65 | 0 | 35 | probably N |
| *Pleuronichthys coenosus* | Pleuronectidae | Pleuronectiformes | demersal | 50 | 46 | 4 | unclear |
| *Pleuronichthys decurrens* | Pleuronectidae | Pleuronectiformes | demersal | 50 | 46 | 4 | unclear |
| *Pleuronichthys ritteri* | Pleuronectidae | Pleuronectiformes | demersal | 50 | 46 | 4 | unclear |
| *Pleuronichthys verticalis* | Pleuronectidae | Pleuronectiformes | demersal | 50 | 46 | 4 | unclear |
| *Psettichthys melanostictus* | Pleuronectidae | Pleuronectiformes | demersal | 93 | 7 | 0 | N |
| *Reinhardtius hippoglossoides* | Pleuronectidae | Pleuronectiformes | demersal | 50 | 46 | 4 | unclear |
| *Oncorhynchus clarkii* | Salmonidae | Salmoniformes | diadromous | 59 | 0 | 41 | probably N |
| *Oncorhynchus gorbuscha* | Salmonidae | Salmoniformes | diadromous | 59 | 0 | 41 | probably N |
| *Oncorhynchus keta* | Salmonidae | Salmoniformes | diadromous | 59 | 0 | 41 | probably N |
| *Oncorhynchus kisutch* | Salmonidae | Salmoniformes | diadromous | 59 | 0 | 41 | probably N |
| *Oncorhynchus mykiss* | Salmonidae | Salmoniformes | diadromous | 61 | 1 | 38 | probably N |
| *Oncorhynchus nerka* | Salmonidae | Salmoniformes | diadromous | 59 | 0 | 41 | probably N |
| *Oncorhynchus tshawytscha* | Salmonidae | Salmoniformes | diadromous | 59 | 0 | 41 | probably N |
| *Brama japonica* | Bramidae | Scombriformes | pelagic | 42 | 52 | 6 | unclear |
| *Taractichthys steindachneri* | Bramidae | Scombriformes | pelagic | 9 | 87 | 4 | S |
| *Icichthys lockingtoni* | Centrolophidae | Scombriformes | pelagic | 47 | 51 | 2 | unclear |
| *Gempylus serpens* | Gempylidae | Scombriformes | pelagic | 1 | 96 | 3 | S |

| | | | | | | | |
|---|---|---|---|---|---|---|---|
| *Lepidocybium flavobrunneum* | Gempylidae | Scombriformes | pelagic | 0 | 92 | 8 | S |
| *Ruvettus pretiosus* | Gempylidae | Scombriformes | demersal | 1 | 94 | 5 | S |
| *Icosteus aenigmaticus* | Icosteidae | Scombriformes | demersal | 91 | 1 | 8 | N |
| *Acanthocybium solandri* | Scombridae | Scombriformes | pelagic | 1 | 93 | 6 | S |
| *Auxis rochei rochei* | Scombridae | Scombriformes | pelagic | 1 | 93 | 6 | S |
| *Auxis thazard thazard* | Scombridae | Scombriformes | pelagic | 1 | 93 | 6 | S |
| *Euthynnus affinis* | Scombridae | Scombriformes | pelagic | 1 | 93 | 6 | S |
| *Euthynnus lineatus* | Scombridae | Scombriformes | pelagic | 1 | 93 | 6 | S |
| *Katsuwonus pelamis* | Scombridae | Scombriformes | pelagic | 1 | 93 | 6 | S |
| *Sarda chiliensis chiliensis* | Scombridae | Scombriformes | pelagic | 2 | 93 | 5 | S |
| *Scomber japonicus* | Scombridae | Scombriformes | pelagic | 40 | 53 | 7 | unclear |
| *Scomberomorus concolor* | Scombridae | Scombriformes | pelagic | 0 | 92 | 8 | S |
| *Scomberomorus sierra* | Scombridae | Scombriformes | pelagic | 0 | 92 | 8 | S |
| *Thunnus alalunga* | Scombridae | Scombriformes | pelagic | 1 | 93 | 6 | S |
| *Thunnus albacares* | Scombridae | Scombriformes | pelagic | 1 | 93 | 6 | S |
| *Thunnus obesus* | Scombridae | Scombriformes | pelagic | 1 | 93 | 6 | S |
| *Thunnus orientalis* | Scombridae | Scombriformes | pelagic | 1 | 93 | 6 | S |
| *Peprilus simillimus* | Stromateidae | Scombriformes | demersal | 43 | 45 | 12 | unclear |
| *Tetragonurus cuvieri* | Tetragonuridae | Scombriformes | pelagic | 5 | 94 | 1 | S |
| *Assurger anzac* | Trichiuridae | Scombriformes | demersal | 0 | 96 | 4 | S |
| *Trichiurus lepturus* | Trichiuridae | Scombriformes | demersal | 0 | 95 | 5 | S |
| *Bagre panamensis* | Ariidae | Siluriformes | demersal | 0 | 80 | 20 | S |
| *Macroramphosus gracilis* | Centriscidae | Syngnathiformes | both demersal and pelagic | 1 | 94 | 5 | S |
| *Fistularia commersonii* | Fistulariidae | Syngnathiformes | demersal | 1 | 92 | 7 | S |
| *Fistularia corneta* | Fistulariidae | Syngnathiformes | demersal | 1 | 92 | 7 | S |
| *Syngnathus auliscus* | Syngnathidae | Syngnathiformes | demersal | 0 | 87 | 13 | S |
| *Syngnathus californiensis* | Syngnathidae | Syngnathiformes | demersal | 0 | 87 | 13 | S |
| *Syngnathus exilis* | Syngnathidae | Syngnathiformes | demersal | 0 | 87 | 13 | S |
| *Syngnathus leptorhynchus* | Syngnathidae | Syngnathiformes | demersal | 0 | 87 | 13 | S |
| *Balistes polylepis* | Balistidae | Tetraodontiformes | demersal | 49 | 47 | 4 | unclear |
| *Xanthichthys mento* | Balistidae | Tetraodontiformes | demersal | 0 | 96 | 4 | S |
| *Chilomycterus reticulatus* | Diodontidae | Tetraodontiformes | demersal | 0 | 97 | 3 | S |
| *Diodon holocanthus* | Diodontidae | Tetraodontiformes | demersal | 0 | 97 | 3 | S |
| *Diodon hystrix* | Diodontidae | Tetraodontiformes | demersal | 0 | 97 | 3 | S |
| *Mola mola* | Molidae | Tetraodontiformes | pelagic | 2 | 94 | 4 | S |
| *Ranzania laevis* | Molidae | Tetraodontiformes | pelagic | 2 | 94 | 4 | S |
| *Lactoria diaphana* | Ostraciidae | Tetraodontiformes | demersal | 0 | 93 | 7 | S |

| | | | | | | | | |
|---|---|---|---|---|---|---|---|---|
| *Lagocephalus lagocephalus lagocephalus* | Tetraodontidae | Tetraodontiformes | pelagic | 0 | 97 | 3 | S | |
| *Sphoeroides annulatus* | Tetraodontidae | Tetraodontiformes | demersal | 0 | 91 | 9 | S | |
| *Sphoeroides lobatus* | Tetraodontidae | Tetraodontiformes | demersal | 0 | 81 | 19 | S | |
| *Allocyttus folletti* | Oreosomatidae | Zeiformes | demersal | 93 | 0 | 7 | N | |
| *Zenopsis nebulosa* | Zeidae | Zeiformes | pelagic | 2 | 1 | 97 | probably N | |
| *Howella brodiei* | Howellidae | Acropomatiformes | pelagic | 49 | 49 | 2 | unclear | Fig. S1 only |
| *Alepocephalus tenebrosus* | Alepocephalidae | Alepocephaliformes | demersal | 40 | 45 | 15 | unclear | Fig. S1 only |
| *Bajacalifornia burragei* | Alepocephalidae | Alepocephaliformes | demersal | 40 | 45 | 15 | unclear | Fig. S1 only |
| *Bathylaco nigricans* | Alepocephalidae | Alepocephaliformes | pelagic | 40 | 45 | 15 | unclear | Fig. S1 only |
| *Talismania bifurcata* | Alepocephalidae | Alepocephaliformes | demersal | 40 | 45 | 15 | unclear | Fig. S1 only |
| *Holtbyrnia latifrons* | Platytroctidae | Alepocephaliformes | pelagic | 40 | 45 | 15 | unclear | Fig. S1 only |
| *Maulisia argipalla* | Platytroctidae | Alepocephaliformes | pelagic | 40 | 45 | 15 | unclear | Fig. S1 only |
| *Mirorictus taningi* | Platytroctidae | Alepocephaliformes | pelagic | 40 | 45 | 15 | unclear | Fig. S1 only |
| *Sagamichthys abei* | Platytroctidae | Alepocephaliformes | pelagic | 40 | 45 | 15 | unclear | Fig. S1 only |
| *Thalassenchelys coheni* | Colocongridae | Anguilliformes | pelagic | 44 | 54 | 2 | unclear | Fig. S1 only |
| *Avocettina infans* | Nemichthyidae | Anguilliformes | pelagic | 27 | 63 | 10 | unclear | Fig. S1 only |
| *Nemichthys scolopaceus* | Nemichthyidae | Anguilliformes | pelagic | 27 | 63 | 10 | unclear | Fig. S1 only |
| *Serrivomer jesperseni* | Serrivomeridae | Anguilliformes | pelagic | 27 | 63 | 10 | unclear | Fig. S1 only |
| *Serrivomer sector* | Serrivomeridae | Anguilliformes | pelagic | 27 | 63 | 10 | unclear | Fig. S1 only |
| *Bathylagus pacificus* | Microstomatidae | Argentiniformes | pelagic | 58 | 39 | 3 | unclear | Fig. S1 only |
| *Lipolagus ochotensis* | Microstomatidae | Argentiniformes | pelagic | 58 | 39 | 3 | unclear | Fig. S1 only |

| | | | | | | | | |
|---|---|---|---|---|---|---|---|---|
| *Pseudobathylagus milleri* | Microstomatidae | Argentiniformes | pelagic | 58 | 39 | 3 | unclear | Fig. S1 only |
| *Macropinna microstoma* | Opisthoproctidae | Argentiniformes | pelagic | 58 | 39 | 3 | unclear | Fig. S1 only |
| *Rhynchohyalus natalensis* | Opisthoproctidae | Argentiniformes | pelagic | 58 | 39 | 3 | unclear | Fig. S1 only |
| *Alepisaurus ferox* | Alepisauridae | Aulopiformes | pelagic | 6 | 91 | 3 | S | Fig. S1 only |
| *Gigantura indica* | Giganturidae | Aulopiformes | pelagic | 1 | 94 | 5 | S | Fig. S1 only |
| *Scopelosaurus harryi* | Notosudidae | Aulopiformes | pelagic | 42 | 54 | 4 | unclear | Fig. S1 only |
| *Anotopterus nikparini* | Paralepididae | Aulopiformes | pelagic | 6 | 91 | 3 | S | Fig. S1 only |
| *Lestidiops pacificus* | Paralepididae | Aulopiformes | pelagic | 6 | 91 | 3 | S | Fig. S1 only |
| *Lestidiops ringens* | Paralepididae | Aulopiformes | pelagic | 6 | 91 | 3 | S | Fig. S1 only |
| *Macroparalepis johnfitchi* | Paralepididae | Aulopiformes | pelagic | 6 | 91 | 3 | S | Fig. S1 only |
| *Magnisudis atlantica* | Paralepididae | Aulopiformes | pelagic | 6 | 91 | 3 | S | Fig. S1 only |
| *Sudis atrox* | Paralepididae | Aulopiformes | pelagic | 6 | 91 | 3 | S | Fig. S1 only |
| *Benthalbella dentata* | Scopelarchidae | Aulopiformes | pelagic | 39 | 53 | 8 | unclear | Fig. S1 only |
| *Benthalbella linguidens* | Scopelarchidae | Aulopiformes | pelagic | 39 | 53 | 8 | unclear | Fig. S1 only |
| *Barbourisia rufa* | Barbourisiidae | Beryciformes | both demersal and pelagic | 51 | 45 | 4 | unclear | Fig. S1 only |
| *Cetostoma regani* | Cetomimidae | Beryciformes | pelagic | 51 | 45 | 4 | unclear | Fig. S1 only |
| *Melamphaes lugubris* | Melamphaidae | Beryciformes | pelagic | 31 | 67 | 2 | unclear | Fig. S1 only |
| *Poromitra crassiceps* | Melamphaidae | Beryciformes | pelagic | 50 | 45 | 5 | unclear | Fig. S1 only |
| *Scopeloberyx robustus* | Melamphaidae | Beryciformes | pelagic | 51 | 44 | 5 | unclear | Fig. S1 only |

| | | | | | | | | |
|---|---|---|---|---|---|---|---|---|
| *Scopelogadus mizolepis bispinosus* | Melamphaidae | Beryciformes | pelagic | 50 | 47 | 3 | unclear | Fig. S1 only |
| *Rondeletia loricata* | Rondeletiidae | Beryciformes | pelagic | 51 | 45 | 4 | unclear | Fig. S1 only |
| *Albatrossia pectoralis* | Macrouridae | Gadiformes | demersal | 97 | 0 | 3 | N | Fig. S1 only |
| *Coelorinchus scaphopsis* | Macrouridae | Gadiformes | demersal | 0 | 95 | 5 | S | Fig. S1 only |
| *Coryphaenoides armatus* | Macrouridae | Gadiformes | both demersal and pelagic | 60 | 31 | 9 | unclear | Fig. S1 only |
| *Coryphaenoides cinereus* | Macrouridae | Gadiformes | demersal | 97 | 0 | 3 | N | Fig. S1 only |
| *Coryphaenoides filifer* | Macrouridae | Gadiformes | demersal | 97 | 0 | 3 | N | Fig. S1 only |
| *Coryphaenoides leptolepis* | Macrouridae | Gadiformes | demersal | 42 | 55 | 3 | unclear | Fig. S1 only |
| *Malacocephalus laevis* | Macrouridae | Gadiformes | both demersal and pelagic | 0 | 94 | 6 | S | Fig. S1 only |
| *Nezumia liolepis* | Macrouridae | Gadiformes | demersal | 51 | 47 | 2 | unclear | Fig. S1 only |
| *Melanonus zugmayeri* | Melanonidae | Gadiformes | pelagic | 44 | 49 | 7 | unclear | Fig. S1 only |
| *Halargyreus johnsonii* | Moridae | Gadiformes | both demersal and pelagic | 89 | 4 | 7 | N | Fig. S1 only |
| *Stylephorus chordatus* | Stylephoridae | Lampridiformes | pelagic | 96 | 0 | 4 | N | Fig. S1 only |
| *Lampris guttatus* | Lampridae | Lampriformes | pelagic | 43 | 51 | 6 | unclear | Fig. S1 only |
| *Regalecus russelii* | Regalecidae | Lampriformes | pelagic | 0 | 95 | 5 | S | Fig. S1 only |
| *Cryptopsaras couesii* | Ceratiidae | Lophiiformes | pelagic | 13 | 83 | 4 | S | Fig. S1 only |
| *Chaunacops coloratus* | Chaunacidae | Lophiiformes | demersal | 5 | 91 | 4 | S | Fig. S1 only |
| *Gigantactis vanhoeffeni* | Gigantactinidae | Lophiiformes | pelagic | 13 | 83 | 4 | S | Fig. S1 only |
| *Melanocetus johnsonii* | Melanocetidae | Lophiiformes | pelagic | 13 | 83 | 4 | S | Fig. S1 only |

| | | | | | | | | |
|---|---|---|---|---|---|---|---|---|
| *Bertella idiomorpha* | Oneirodidae | Lophiiformes | pelagic | 13 | 83 | 4 | S | Fig. S1 only |
| *Oneirodes acanthias* | Oneirodidae | Lophiiformes | pelagic | 38 | 55 | 7 | unclear | Fig. S1 only |
| *Oneirodes thompsoni* | Oneirodidae | Lophiiformes | pelagic | 41 | 52 | 7 | unclear | Fig. S1 only |
| *Benthosema panamense* | Myctophidae | Myctophiformes | pelagic | 0 | 96 | 4 | S | Fig. S1 only |
| *Ceratoscopelus townsendi* | Myctophidae | Myctophiformes | pelagic | 52 | 45 | 3 | unclear | Fig. S1 only |
| *Diaphus anderseni* | Myctophidae | Myctophiformes | pelagic | 0 | 98 | 2 | S | Fig. S1 only |
| *Diaphus theta* | Myctophidae | Myctophiformes | pelagic | 52 | 45 | 3 | unclear | Fig. S1 only |
| *Diogenichthys laternatus* | Myctophidae | Myctophiformes | pelagic | 0 | 96 | 4 | S | Fig. S1 only |
| *Electrona risso* | Myctophidae | Myctophiformes | pelagic | 45 | 49 | 6 | unclear | Fig. S1 only |
| *Gonichthys tenuiculus* | Myctophidae | Myctophiformes | pelagic | 0 | 94 | 6 | S | Fig. S1 only |
| *Hygophum proximum* | Myctophidae | Myctophiformes | pelagic | 0 | 96 | 4 | S | Fig. S1 only |
| *Hygophum reinhardtii* | Myctophidae | Myctophiformes | pelagic | 0 | 96 | 4 | S | Fig. S1 only |
| *Lampadena urophaos urophaos* | Myctophidae | Myctophiformes | pelagic | 52 | 45 | 3 | unclear | Fig. S1 only |
| *Lampanyctus tenuiformis* | Myctophidae | Myctophiformes | pelagic | 2 | 87 | 11 | S | Fig. S1 only |
| *Myctophum nitidulum* | Myctophidae | Myctophiformes | pelagic | 4 | 92 | 4 | S | Fig. S1 only |
| *Nannobrachium regale* | Myctophidae | Myctophiformes | pelagic | 2 | 87 | 11 | S | Fig. S1 only |
| *Nannobrachium ritteri* | Myctophidae | Myctophiformes | pelagic | 2 | 87 | 11 | S | Fig. S1 only |
| *Notoscopelus resplendens* | Myctophidae | Myctophiformes | pelagic | 4 | 93 | 3 | S | Fig. S1 only |
| *Parvilux ingens* | Myctophidae | Myctophiformes | pelagic | 52 | 45 | 3 | unclear | Fig. S1 only |

| | | | | | | | | |
|---|---|---|---|---|---|---|---|---|
| *Protomyctophum crockeri* | Myctophidae | Myctophiformes | pelagic | 87 | 3 | 10 | N | Fig. S1 only |
| *Protomyctophum thompsoni* | Myctophidae | Myctophiformes | pelagic | 87 | 3 | 10 | N | Fig. S1 only |
| *Stenobrachius leucopsarus* | Myctophidae | Myctophiformes | pelagic | 52 | 45 | 3 | unclear | Fig. S1 only |
| *Stenobrachius nannochir* | Myctophidae | Myctophiformes | pelagic | 52 | 45 | 3 | unclear | Fig. S1 only |
| *Symbolophorus californiensis* | Myctophidae | Myctophiformes | pelagic | 0 | 96 | 4 | S | Fig. S1 only |
| *Symbolophorus evermanni* | Myctophidae | Myctophiformes | pelagic | 0 | 96 | 4 | S | Fig. S1 only |
| *Taaningichthys bathyphilus* | Myctophidae | Myctophiformes | pelagic | 52 | 45 | 3 | unclear | Fig. S1 only |
| *Tarletonbeania crenularis* | Myctophidae | Myctophiformes | pelagic | 7 | 89 | 4 | S | Fig. S1 only |
| *Triphoturus mexicanus* | Myctophidae | Myctophiformes | pelagic | 52 | 45 | 3 | unclear | Fig. S1 only |
| *Neoscopelus macrolepidotus* | Neoscopelidae | Myctophiformes | demersal | 69 | 26 | 5 | unclear | Fig. S1 only |
| *Scopelengys tristis* | Neoscopelidae | Myctophiformes | pelagic | 61 | 37 | 2 | unclear | Fig. S1 only |
| *Cataetyx rubrirostris* | Bythitidae | Ophidiiformes | demersal | 50 | 41 | 9 | unclear | Fig. S1 only |
| *Lamprogrammus niger* | Ophidiidae | Ophidiiformes | both demersal and pelagic | 39 | 56 | 5 | unclear | Fig. S1 only |
| *Acantholiparis opercularis* | Liparidae | Perciformes/Cottoidei | demersal | 96 | 1 | 3 | N | Fig. S1 only |
| *Careproctus colletti* | Liparidae | Perciformes/Cottoidei | demersal | 96 | 1 | 3 | N | Fig. S1 only |
| *Careproctus cypselurus* | Liparidae | Perciformes/Cottoidei | demersal | 96 | 1 | 3 | N | Fig. S1 only |
| *Careproctus gilberti* | Liparidae | Perciformes/Cottoidei | demersal | 96 | 1 | 3 | N | Fig. S1 only |
| *Elassodiscus caudatus* | Liparidae | Perciformes/Cottoidei | demersal | 96 | 1 | 3 | N | Fig. S1 only |
| *Nectoliparis pelagicus* | Liparidae | Perciformes/Cottoidei | pelagic | 96 | 1 | 3 | N | Fig. S1 only |

| | | | | | | | | |
|---|---|---|---|---|---|---|---|---|
| *Paraliparis cephalus* | Liparidae | Perciformes/Cottoidei | demersal | 95 | 2 | 3 | N | Fig. S1 only |
| *Paraliparis dactylosus* | Liparidae | Perciformes/Cottoidei | demersal | 96 | 1 | 3 | N | Fig. S1 only |
| *Paraliparis pectoralis* | Liparidae | Perciformes/Cottoidei | demersal | 96 | 1 | 3 | N | Fig. S1 only |
| *Paraliparis rosaceus* | Liparidae | Perciformes/Cottoidei | demersal | 96 | 1 | 3 | N | Fig. S1 only |
| *Rhinoliparis attenuatus* | Liparidae | Perciformes/Cottoidei | pelagic | 96 | 1 | 3 | N | Fig. S1 only |
| *Rhinoliparis barbulifer* | Liparidae | Perciformes/Cottoidei | demersal | 96 | 1 | 3 | N | Fig. S1 only |
| *Psychrolutes phrictus* | Psychrolutidae | Perciformes/Cottoidei | demersal | 100 | 0 | 0 | N | Fig. S1 only |
| *Lycenchelys camchatica* | Zoarcidae | Perciformes/Zoarcoidei | demersal | 73 | 0 | 27 | N | Fig. S1 only |
| *Lycodapus endemoscotus* | Zoarcidae | Perciformes/Zoarcoidei | pelagic | 73 | 0 | 27 | N | Fig. S1 only |
| *Lycodapus pachysoma* | Zoarcidae | Perciformes/Zoarcoidei | pelagic | 73 | 0 | 27 | N | Fig. S1 only |
| *Pachycara gymninium* | Zoarcidae | Perciformes/Zoarcoidei | demersal | 73 | 0 | 27 | N | Fig. S1 only |
| *Pachycara lepinium* | Zoarcidae | Perciformes/Zoarcoidei | demersal | 73 | 0 | 27 | N | Fig. S1 only |
| *Atheresthes evermanni* | Pleuronectidae | Pleuronectiformes | demersal | 55 | 41 | 4 | unclear | Fig. S1 only |
| *Clidoderma asperrimum* | Pleuronectidae | Pleuronectiformes | demersal | 50 | 46 | 4 | unclear | Fig. S1 only |
| *Embassichthys bathybius* | Pleuronectidae | Pleuronectiformes | demersal | 50 | 46 | 4 | unclear | Fig. S1 only |
| *Eurypharynx pelecanoides* | Eurypharyngidae | Saccopharyngiformes | pelagic | 50 | 45 | 5 | unclear | Fig. S1 only |
| *Taractes asper* | Bramidae | Scombriformes | pelagic | 12 | 84 | 4 | S | Fig. S1 only |
| *Chiasmodon niger* | Chiasmodontidae | Scombriformes | pelagic | 41 | 52 | 7 | unclear | Fig. S1 only |
| *Kali indica* | Chiasmodontidae | Scombriformes | pelagic | 6 | 83 | 11 | S | Fig. S1 only |

| | | | | | | | | |
|---|---|---|---|---|---|---|---|---|
| *Diplospinus multistriatus* | Gempylidae | Scombriformes | pelagic | 0 | 92 | 8 | S | Fig. S1 only |
| *Cubiceps baxteri* | Nomeidae | Scombriformes | pelagic | 0 | 94 | 6 | S | Fig. S1 only |
| *Cubiceps capensis* | Nomeidae | Scombriformes | pelagic | 0 | 97 | 3 | S | Fig. S1 only |
| *Psenes pellucidus* | Nomeidae | Scombriformes | pelagic | 0 | 97 | 3 | S | Fig. S1 only |
| *Aphanopus arigato* | Trichiuridae | Scombriformes | demersal | 51 | 40 | 9 | unclear | Fig. S1 only |
| *Chauliodus macouni* | Chauliodontidae | Stomiiformes | pelagic | 67 | 31 | 2 | unclear | Fig. S1 only |
| *Cyclothone acclinidens* | Gonostomatidae | Stomiiformes | pelagic | 8 | 87 | 5 | S | Fig. S1 only |
| *Cyclothone alba* | Gonostomatidae | Stomiiformes | pelagic | 8 | 87 | 5 | S | Fig. S1 only |
| *Cyclothone atraria* | Gonostomatidae | Stomiiformes | pelagic | 8 | 87 | 5 | S | Fig. S1 only |
| *Cyclothone microdon* | Gonostomatidae | Stomiiformes | pelagic | 8 | 87 | 5 | S | Fig. S1 only |
| *Cyclothone pallida* | Gonostomatidae | Stomiiformes | pelagic | 8 | 87 | 5 | S | Fig. S1 only |
| *Cyclothone pseudopallida* | Gonostomatidae | Stomiiformes | pelagic | 8 | 87 | 5 | S | Fig. S1 only |
| *Cyclothone signata* | Gonostomatidae | Stomiiformes | pelagic | 8 | 87 | 5 | S | Fig. S1 only |
| *Diplophos taenia* | Gonostomatidae | Stomiiformes | pelagic | 5 | 90 | 5 | S | Fig. S1 only |
| *Gonostoma atlanticum* | Gonostomatidae | Stomiiformes | pelagic | 8 | 87 | 5 | S | Fig. S1 only |
| *Sigmops gracilis* | Gonostomatidae | Stomiiformes | pelagic | 8 | 87 | 5 | S | Fig. S1 only |
| *Idiacanthus antrostomus* | Idiacanthidae | Stomiiformes | pelagic | 36 | 60 | 4 | unclear | Fig. S1 only |
| *Aristostomias scintillans* | Malacosteidae | Stomiiformes | pelagic | 41 | 54 | 5 | unclear | Fig. S1 only |
| *Photonectes margarita* | Melanostomiidae | Stomiiformes | pelagic | 13 | 84 | 3 | S | Fig. S1 only |

| | | | | | | | | |
|---|---|---|---|---|---|---|---|---|
| *Tactostoma macropus* | Melanostomiidae | Stomiiformes | pelagic | 37 | 59 | 4 | unclear | Fig. S1 only |
| *Vinciguerria nimbaria* | Phosichthyidae | Stomiiformes | pelagic | 7 | 88 | 5 | S | Fig. S1 only |
| *Argyropelecus aculeatus* | Sternoptychidae | Stomiiformes | pelagic | 31 | 65 | 4 | unclear | Fig. S1 only |
| *Argyropelecus affinis* | Sternoptychidae | Stomiiformes | pelagic | 31 | 65 | 4 | unclear | Fig. S1 only |
| *Argyropelecus hemigymnus* | Sternoptychidae | Stomiiformes | pelagic | 31 | 65 | 4 | unclear | Fig. S1 only |
| *Argyropelecus lychnus* | Sternoptychidae | Stomiiformes | pelagic | 31 | 65 | 4 | unclear | Fig. S1 only |
| *Argyropelecus sladeni* | Sternoptychidae | Stomiiformes | pelagic | 31 | 65 | 4 | unclear | Fig. S1 only |
| *Danaphos oculatus* | Sternoptychidae | Stomiiformes | pelagic | 9 | 91 | 0 | S | Fig. S1 only |
| *Sternoptyx diaphana* | Sternoptychidae | Stomiiformes | pelagic | 7 | 88 | 5 | S | Fig. S1 only |
| *Sternoptyx obscura* | Sternoptychidae | Stomiiformes | pelagic | 7 | 88 | 5 | S | Fig. S1 only |
| *Sternoptyx pseudobscura* | Sternoptychidae | Stomiiformes | pelagic | 7 | 88 | 5 | S | Fig. S1 only |
| *Borostomias panamensis* | Stomiidae | Stomiiformes | pelagic | 52 | 44 | 4 | unclear | Fig. S1 only |
| *Chauliodus sloani* | Stomiidae | Stomiiformes | pelagic | 73 | 25 | 2 | N | Fig. S1 only |
| *Flagellostomias boureei* | Stomiidae | Stomiiformes | pelagic | 0 | 96 | 4 | S | Fig. S1 only |
| *Idiacanthus fasciola* | Stomiidae | Stomiiformes | pelagic | 36 | 60 | 4 | unclear | Fig. S1 only |
| *Malacosteus niger* | Stomiidae | Stomiiformes | pelagic | 41 | 54 | 5 | unclear | Fig. S1 only |
| *Neonesthes capensis* | Stomiidae | Stomiiformes | pelagic | 18 | 22 | 60 | unclear | Fig. S1 only |
| *Pachystomias microdon* | Stomiidae | Stomiiformes | pelagic | 41 | 54 | 5 | unclear | Fig. S1 only |
| *Rhadinesthes decimus* | Stomiidae | Stomiiformes | pelagic | 45 | 49 | 6 | unclear | Fig. S1 only |

| | | | | | | | | |
|---|---|---|---|---|---|---|---|---|
| *Stomias atriventer* | Stomiidae | Stomiiformes | pelagic | 41 | 52 | 7 | unclear | Fig. S1 only |
| *Anoplogaster cornuta* | Anoplogastridae | Trachichthyiformes | pelagic | 53 | 43 | 4 | unclear | Fig. S1 only |
| *Diretmus argenteus* | Diretmidae | Trachichthyiformes | pelagic | 0 | 91 | 9 | S | Fig. S1 only |

**Table S5:** Region-of-origin for California cartilaginous fishes as inferred from BioGeoBEARS analyses.

| species | order | family | habitat | # maps inferring northern origin | # maps inferring southern origin | # maps indet. | region of origin coded in plots | notes |
|---|---|---|---|---|---|---|---|---|
| *Carcharhinus brachyurus* | Carcharhiniformes | Carcharhinidae | pelagic | 0 | 95 | 5 | S | |
| *Carcharhinus leucas* | Carcharhiniformes | Carcharhinidae | pelagic | 0 | 96 | 4 | S | |
| *Carcharhinus obscurus* | Carcharhiniformes | Carcharhinidae | pelagic | 0 | 98 | 2 | S | |
| *Nasolamia velox* | Carcharhiniformes | Carcharhinidae | demersal | 0 | 95 | 5 | S | |
| *Prionace glauca* | Carcharhiniformes | Carcharhinidae | pelagic | 1 | 97 | 2 | S | |
| *Rhizoprionodon longurio* | Carcharhiniformes | Carcharhinidae | demersal | 0 | 94 | 6 | S | |
| *Carcharodon carcharias* | Carcharhiniformes | Lamnidae | pelagic | 10 | 88 | 2 | S | |
| *Apristurus brunneus* | Carcharhiniformes | Pentanchidae | both demersal and pelagic | 44 | 50 | 6 | unclear | |
| *Apristurus kampae* | Carcharhiniformes | Pentanchidae | demersal | 44 | 53 | 3 | unclear | |
| *Parmaturus xaniurus* | Carcharhiniformes | Pentanchidae | demersal | 53 | 41 | 6 | unclear | |
| *Cephaloscyllium ventriosum* | Carcharhiniformes | Scyliorhinidae | demersal | 0 | 94 | 6 | S | |
| *Sphyrna lewini* | Carcharhiniformes | Sphyrnidae | pelagic | 0 | 99 | 1 | S | |
| *Galeorhinus galeus* | Carcharhiniformes | Triakidae | pelagic | 43 | 49 | 8 | unclear | |
| *Mustelus californicus* | Carcharhiniformes | Triakidae | demersal | 0 | 97 | 3 | S | |
| *Mustelus henlei* | Carcharhiniformes | Triakidae | demersal | 42 | 45 | 13 | unclear | |
| *Mustelus lunulatus* | Carcharhiniformes | Triakidae | demersal | 0 | 96 | 4 | S | |
| *Triakis semifasciata* | Carcharhiniformes | Triakidae | demersal | 32 | 42 | 26 | unclear | |
| *Hydrolagus colliei* | Chimaeriformes | Chimaeridae | demersal | 51 | 40 | 9 | unclear | |
| *Hydrolagus melanophasma* | Chimaeriformes | Chimaeridae | demersal | 1 | 92 | 7 | S | |
| *Echinorhinus cookei* | Echinorhiniformes | Echinorhinidae | demersal | 49 | 46 | 5 | unclear | |
| *Heterodontus francisci* | Heterodontiformes | Heterodontidae | demersal | 1 | 96 | 3 | S | |
| *Chlamydoselachus anguineus* | Hexanchiformes | Chlamydoselachidae | demersal | 28 | 69 | 3 | unclear | |
| *Hexanchus griseus* | Hexanchiformes | Hexanchidae | demersal | 28 | 69 | 3 | unclear | |
| *Notorynchus cepedianus* | Hexanchiformes | Hexanchidae | demersal | 28 | 69 | 3 | unclear | |
| *Alopias superciliosus* | Lamniformes | Alopiidae | pelagic | 10 | 88 | 2 | S | |
| *Alopias vulpinus* | Lamniformes | Alopiidae | pelagic | 10 | 88 | 2 | S | |
| *Cetorhinus maximus* | Lamniformes | Cetorhinidae | pelagic | 10 | 88 | 2 | S | |

| | | | | | | | | |
|---|---|---|---|---|---|---|---|---|
| *Isurus oxyrinchus* | Lamniformes | Lamnidae | pelagic | 10 | 88 | 2 | S | |
| *Lamna ditropis* | Lamniformes | Lamnidae | pelagic | 10 | 88 | 2 | S | |
| *Dasyatis dipterura* | Myliobatiformes | Dasyatidae | demersal | 0 | 91 | 9 | S | |
| *Pteroplatytrygon violacea* | Myliobatiformes | Dasyatidae | pelagic | 41 | 43 | 16 | unclear | |
| *Gymnura marmorata* | Myliobatiformes | Gymnuridae | demersal | 0 | 93 | 7 | S | |
| *Manta birostris* | Myliobatiformes | Mobulidae | pelagic | 0 | 100 | 0 | S | |
| *Myliobatis californicus* | Myliobatiformes | Myliobatidae | demersal | 38 | 53 | 9 | unclear | |
| *Urobatis halleri* | Myliobatiformes | Urotrygonidae | demersal | 0 | 100 | 0 | S | |
| *Rhincodon typus* | Orectolobiformes | Rhincodontidae | pelagic | 1 | 87 | 12 | S | |
| *Bathyraja abyssicola* | Rajiformes | Arhynchobatidae | demersal | 98 | 1 | 1 | N | |
| *Bathyraja aleutica* | Rajiformes | Arhynchobatidae | demersal | 98 | 1 | 1 | N | |
| *Amblyraja hyperborea* | Rajiformes | Rajidae | demersal | 53 | 44 | 3 | unclear | |
| *Beringraja binoculata* | Rajiformes | Rajidae | demersal | 52 | 44 | 4 | unclear | |
| *Raja inornata* | Rajiformes | Rajidae | demersal | 52 | 44 | 4 | unclear | |
| *Raja rhina* | Rajiformes | Rajidae | demersal | 52 | 44 | 4 | unclear | |
| *Raja stellulata* | Rajiformes | Rajidae | demersal | 52 | 44 | 4 | unclear | |
| *Rhinobatos productus* | Rhinopristiformes | Rhinobatidae | demersal | 0 | 98 | 2 | S | |
| *Zapteryx exasperata* | Rhinopristiformes | Trygonorrhinidae | demersal | 1 | 94 | 5 | S | |
| *Euprotomicrus bispinatus* | Squaliformes | Dalatiidae | pelagic | 0 | 99 | 1 | S | |
| *Isistius brasiliensis* | Squaliformes | Dalatiidae | pelagic | 0 | 97 | 3 | S | |
| *Somniosus pacificus* | Squaliformes | Somniosidae | demersal | 47 | 48 | 5 | unclear | |
| *Zameus squamulosus* | Squaliformes | Somniosidae | demersal | 0 | 95 | 5 | S | |
| *Squalus acanthias* | Squaliformes | Squalidae | demersal | 40 | 28 | 32 | unclear | |
| *Squatina californica* | Squatiniformes | Squatinidae | demersal | 43 | 53 | 4 | unclear | |
| *Platyrhinoidis triseriata* | Torpediniformes | Platyrhinidae | demersal | 0 | 96 | 4 | S | |
| *Torpedo californica* | Torpediniformes | Torpedinidae | demersal | 47 | 48 | 5 | unclear | |
| *Harriotta raleighana* | Chimaeriformes | Rhinochimaeridae | demersal | 0 | 91 | 9 | S | Fig. S1 only |
